# Cancer-associated HIF-2α impacts trunk neural crest stemness

**DOI:** 10.1101/2020.01.22.915199

**Authors:** Sofie Mohlin, Camilla U. Persson, Elina Fredlund, Emanuela Monni, Jessica M. Lindvall, Zaal Kokaia, Emma Hammarlund, Marianne E. Bronner

## Abstract

The neural crest is a stem cell population that gives rise to sympathetic ganglia, the cell type of origin of neuroblastoma. Hypoxia Inducible Factor (HIF)-2α is associated with high risk neuroblastoma, however, little is known about its role in normal neural crest development. To address this important question, here we show that HIF-2α is expressed in trunk neural crest cells of human, murine and avian embryos. Modulating HIF-2*α in vivo* not only causes developmental delays but also induces proliferation and stemness of neural crest cells while altering the number of cells migrating ventrally to sympathoadrenal sites. Transcriptome changes after loss of HIF-2α reflect the *in vivo* phenotype. The results suggest that expression levels of HIF-2α must be strictly controlled and abnormal levels increase stemness and may promote metastasis. Our findings help elucidate the role of HIF-2α during normal development with implications also in tumor initiation at the onset of neuroblastoma.

## Introduction

The neural crest is a multipotent stem cell population that is unique to vertebrate embryos. Originating from the ectodermal germ layer, premigratory neural crest cells arise in the dorsal neural tube during neurulation and are characterized by expression of transcription factors like *FOXD3*, *TFAP2* and *SOXE* (Khudyakov & Bronner-Fraser, 2009). Neural crest cells subsequently undergo an epithelial-to-mesenchymal transition (EMT) to delaminate from the neuroepithelium, then migrate extensively throughout the embryo, populating distant sites. Upon reaching their final destinations, neural crest cells form a large variety of cell types, as diverse as elements of the craniofacial skeleton, melanocytes of the skin, adrenal chromaffin cells and sympathetic neurons and glia (Ayer-Le Lievre & Le Douarin, 1982; Bittencourt, da Costa, Calloni, Alvarez-Silva, & Trentin, 2013; Bronner-Fraser & Fraser, 1988; Vega-Lopez, Cerrizuela, Tribulo, & Aybar, 2018).

The stem cell properties and migratory nature of the neural crest are highly reminiscent of tumor cells. Indeed, many of the genes involved in neural crest EMT are redeployed in metastatic cancers including many types of neural crest-derived cancers. Thus, neural crest cells represent an excellent model for studying the origin of neural crest-derived tumors including pediatric neuroblastoma, a tumor of infancy responsible for 15% of all cancer-related deaths in children (Maris, 2010). Neuroblastoma patients are very young, with some tumors detected in newborns. It is well accepted that neuroblastoma derives from sympathetic neuroblasts that originate from trunk neural crest cells (De Preter et al., 2006; Hoehner et al., 1996).

High risk neuroblastoma correlates with the presence of cells in perivascular niches (Pietras et al., 2008) that express high levels of Hypoxia Inducible Factor (HIF)-2α together with numerous neural crest markers (Holmquist-Mengelbier et al., 2006; Pietras et al., 2008; Pietras et al., 2009). Under normal conditions, HIF-2α is stabilized at low oxygen levels and responds to hypoxia by initiating a transcriptional program for cellular adaptation to changes in metabolic demand. In neuroblastoma, however, HIF-2α becomes abnormally stabilized at physiological oxygen tensions (∼5% O_2_) (Holmquist-Mengelbier et al., 2006). This, together with the presence of neural crest markers in neuroblastoma tumors, raises the intriguing possibility that HIF-2α expressing neural crest cells in the early embryo might reflect the cell type of origin in tumor initiation.

Here, we explore this possibility by examining the role of HIF-2α, encoded by the gene *EPAS1,* during normal neural crest development and possible correlations with neuroblastoma. We show that HIF-2α is expressed in migrating trunk neural crest and sympathetic neuroblasts in human, murine and avian embryos. RNA sequencing of trunk neural crest cells with dysregulated HIF-2α levels demonstrates a shift in the global transcriptional program, resulting in enrichment in genes associated with processes connected to tumor morphology, invasion, EMT and arrested embryo growth. Perturbation experiments in chick embryos *in vivo* result in a delay in embryonic growth, altered expression of trunk neural crest genes, and disrupted trunk neural crest cell migration. Consistent with this, *in vitro* crestospheres display increased proliferation and self-renewal capacity. The results suggest that expression levels of HIF-2α must be tightly regulated. These findings enhance our understanding of how genes dysregulated in normal development may result in onset of neuroblastoma.

## Results

### HIF-2α is expressed in migratory trunk neural crest cells in chick embryos

The presence of neuroblastoma cells expressing hypoxia inducible factor (HIF)-2α in perivascular tumor niches indicates poor prognosis in this tumor form. That these cells express stem cell- and neural crest associated proteins raises the intriguing possibility that they may constitute a tumor-initiating subpopulation of cells that resembles embryonic neural crest cells. HIF-2α is a transcription factor that localizes to the nucleus but also is found in the cytoplasm (Holmquist-Mengelbier et al., 2006; Mohlin, Hamidian, & Pahlman, 2013), though its role in the cytoplasm remains unknown. Consistent with this dual localization, Western blots of stage HH18 wild type chick embryos revealed expression of HIF-2α in both the nuclear and cytoplasmic fractions (**Figure 1A**), similar to what has been observed in oxygenated neuroblastoma cells (Holmquist-Mengelbier et al., 2006).

**Fig. 1.**
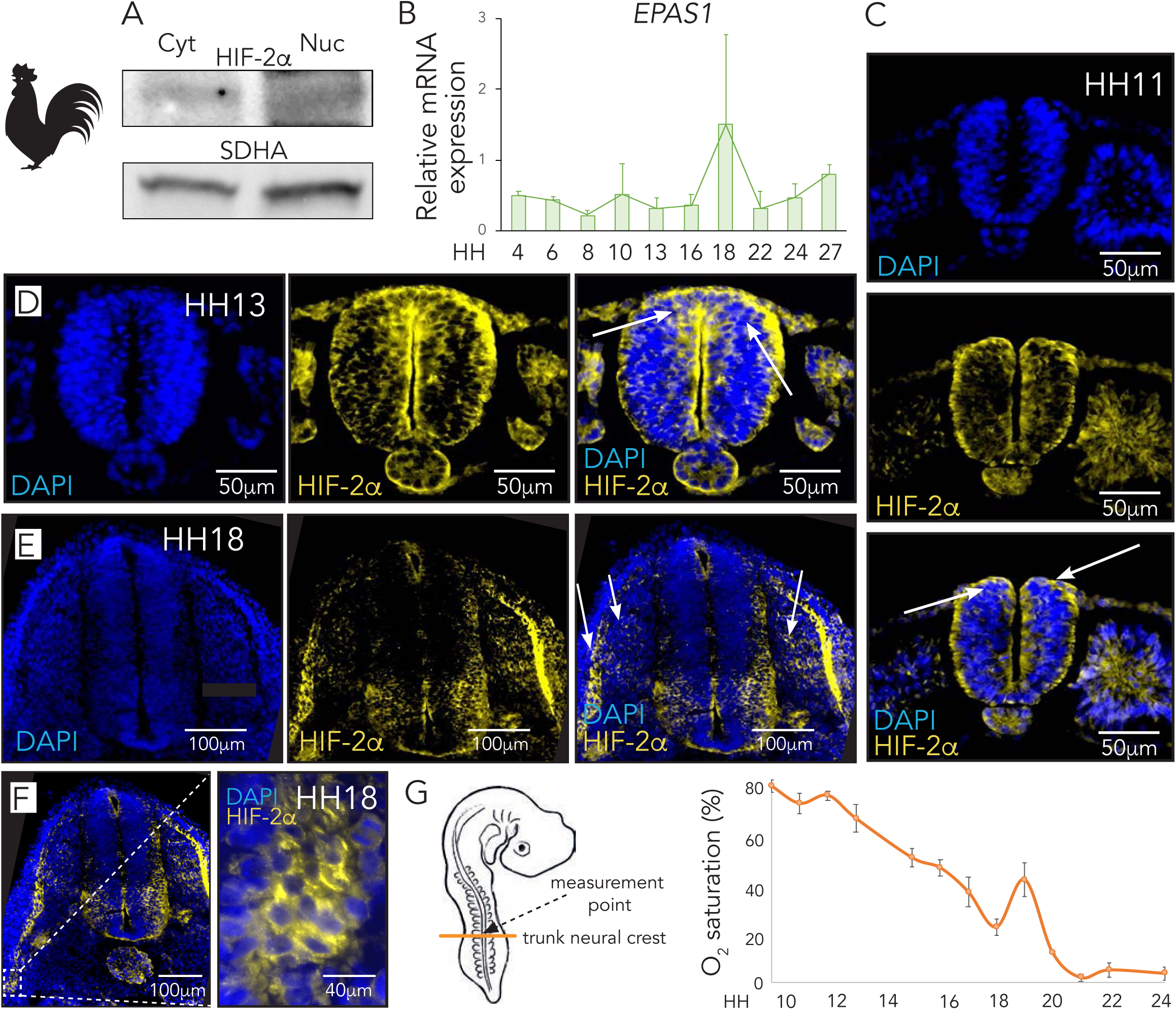
HIF-2α is expressed in trunk neural crest cells. **A.** Western blot of fractionated wild type HH18 chick embryos show HIF-2α protein expression in cytoplasmic and nuclear compartments (cf. panel **C-F**). Blot shown is a representative of multiple experiments. SDHA was used as loading control. **B.** Relative mRNA expression over developmental time (HH4 to HH27) in whole wild type chick embryos. *EPAS1* expression was measured using qRT-PCR and is presented as mean of n=2 biological replicates. Error bars represent SEM. **C-D.** Immunostaining of HIF-2α in sections from trunk axial level of wild type chick embryos at premigratory HH11 **(C)** and HH13 **(D)** stages. Arrows denote scattered HIF-2α positive cells within the dorsal neural tube. **E.** Immunostaining of HIF-2α in sections from trunk axial level of wild type chick embryos at migratory HH18 stage. Arrows denote ventrally migrating HIF-2α positive cells. **F.** A different section from embryo in **(E)** with magnification (dashed square). **G.** Oxygen saturation (%) in the trunk of chick embryos during development measured *ex ovo* using microsensor technique. Error bars represent SEM.

As a first step in exploring the role of HIF-2α in the embryo, we examined its spatiotemporal expression during normal neural crest development. To this end, RNA was extracted from whole chick embryos from stages HH4 to HH27, reflecting stages from gastrulation to mid-gestation. The results revealed continuous expression of HIF-2α (encoded by the gene *EPAS1*) over the time course analyzed, with a peak at HH18 which reflects the time of active trunk neural crest migration (**Figure 1B**). Next, we performed immunocytochemistry with an antibody against HIF-2α in transverse sections through the trunk region of stage HH11, HH13 and HH18 embryos. We detected HIF-2α protein in scattered neural crest cells within the neural tube of HH11 and HH13 embryos, stages when trunk neural crest cells are still premigratory (**Figure 1C-D**, respectively). We further detected HIF-2α in trunk neural crest cells that had delaminated from the neural tube and initiated migration (**Figure 1E-F**). Possible non-specific binding by the primary antibody was ruled out by secondary antibody only staining (**Supplementary Figure S1A**).

HIF-2α is canonically induced at low oxygen levels. To understand variations in oxygen consumption during the developmental stages of interest, we measured O_2_ saturation in real-time in the developing chick embryo utilizing a microsensor technique (**Figure 1G**). Within the trunk neural tube, oxygen saturation starts out high (up to 85% ± 5 SEM O_2_ saturation) at premigratory to migratory stages of neural crest development (HH10 – HH16) and gradually decreases (**Figure 1G**). At the time when the majority of trunk neural crest cells have delaminated from the tube (HH18), oxygen saturation is low (23% ± 10 SEM O_2_ saturation), only to rise at later time points (**Figure 1G**). Together with the expression data above, the results suggest that HIF-2α is independent of oxygen availability in the developing embryo (**Figure 1C-G** and **Figure 2**).

**Fig. 2.**
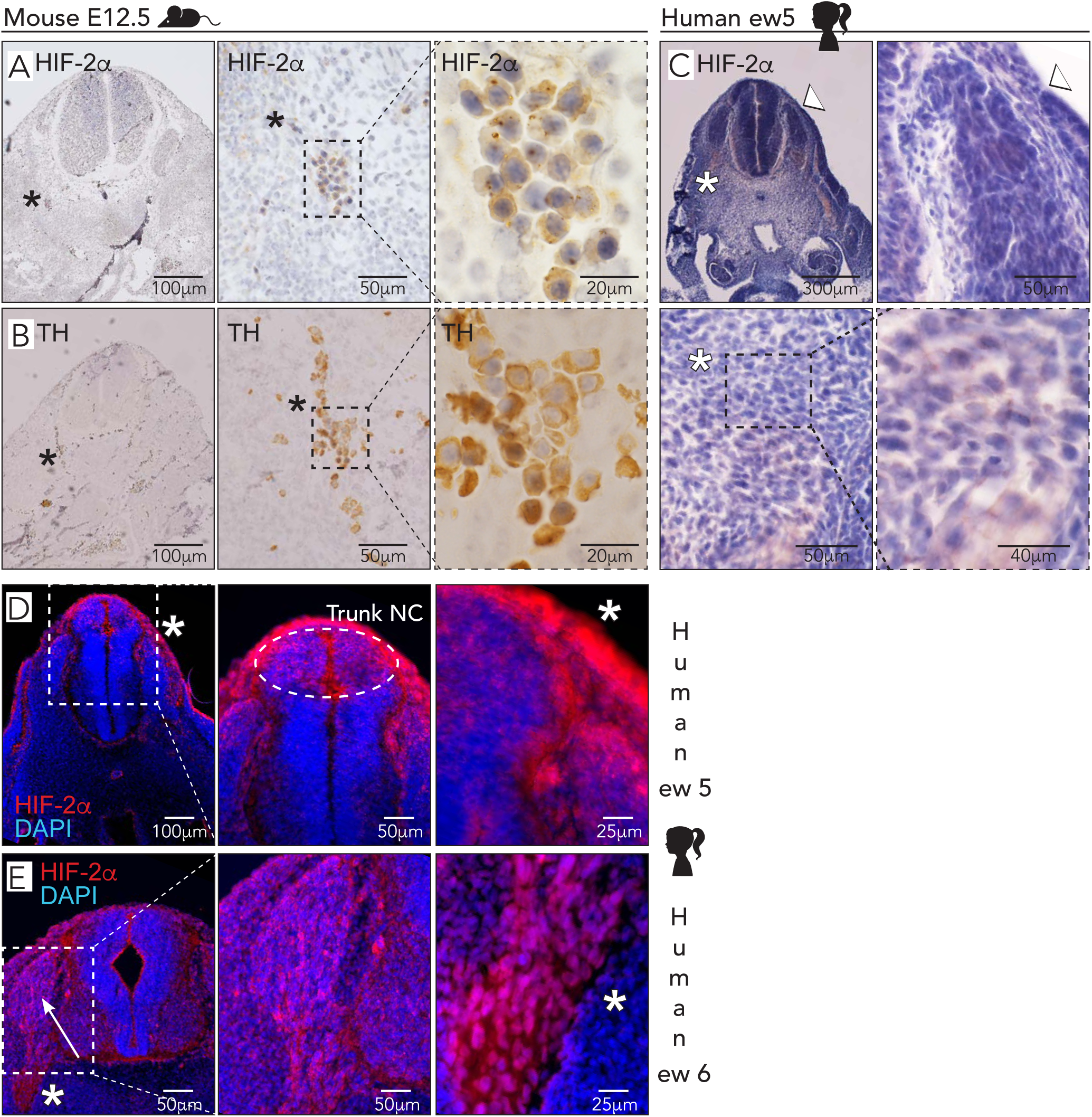
HIF-2α is expressed in human and mouse trunk neural crest cells. **A.** Immunohistochemical staining of HIF-2α in sections from a mouse embryo at embryonic day E12.5. **B.** Immunohistochemical staining of TH in adjacent section to **(A)** to locate sympathetic ganglia. **A-B.** Asterisk in left panels locate HIF-2α^+^ and TH^+^ cells within sympathetic ganglia. Asterisk also indicate magnified area in middle panels and dashed square indicates magnification area in right panels. **C.** Immunohistochemical staining of HIF-2α in sections from trunk axial level of a human embryo at embryonic week ew5. Arrowhead represents magnification in upper right panel. Asterisk represents magnification in lower left panel and dashed square represent magnified area in lower right panel. **(A-C)** Sections are counterstained with hematoxylin to visualize tissue structure and nuclei. **D-E.** Immunostaining of HIF-2α in sections from trunk axial level of human embryos at embryonic week ew5 **(D)** and embryonic week ew6 **(E)**. Arrow denotes HIF-2α positive migrating cells. ew, embryonic week; NC, neural crest. DAPI was used to stain nuclei.

### HIF-2α is expressed in sympathetic neuroblasts in human and mouse embryos

*EPAS1* knockout mice have severe abnormalities in the sympathetic nervous system (Tian, Hammer, Matsumoto, Russell, & McKnight, 1998); consistent with this, there is some, albeit limited, data suggesting that HIF-2α is expressed in sympathetic chain ganglia up to murine day E11.5 (corresponding to human embryonic week 5). Moreover, mice lacking *PHD3* (HIF prolyl hydroxylase), a gene critical for regulation of HIF-2α, display reduced sympathetic nervous system (SNS) function that is rescued by crossing these mutants with EPAS1^+/-^ mice (Bishop et al., 2008).

We have previously shown that HIF-2α is expressed in sympathetic ganglia of human embryos at embryonic week 6.5 (∼E12.5 in mice) but that expression is lost in these cells at later stages (fetal week 8) (Mohlin et al., 2013). Here, we confirmed expression of HIF-2α in sympathetic ganglia in mouse embryos at E12.5 by staining adjacent sections with HIF-2α (**Figure 2A**) and TH (**Figure 2B**) antibodies, with the latter indicating the location of sympathetic ganglia. Demonstrating antibody specificity, HIF-2α expression was only observed in conventional neuroblastoma SK-N-BE(2)c cells cultured at hypoxia (1% O_2_) but not normoxia (21% O_2_) (**Supplementary Figure S1B**). In sections, we detected HIF-2α positive cells specifically in the dorsal neural tube, as well as in early neural crest migratory streams in sections through the trunk region of a human embryo of embryonic week ew5 (Carnegie stage 13; **Figure 2C-D**). In contrast, there were virtually no HIF-2α positive cells left within the neural tube at embryonic week ew6 (Carnegie stage 16). Rather, positive cells could be detected migrating along the ventral pathway followed by sympathoadrenal precursors (**Figure 2E**). Consistent with our biochemical analysis in the chick (**Figure 1A**), human HIF-2α expression was noted in both the nucleus and cytoplasm (**Figure 2E**).

### Knockdown of HIF-2α delays embryogenesis, alters gene expression and affects cell numbers along the ventral neural crest migratory pathway

To examine the role of HIF-2*α in vivo*, we performed loss-of-function experiments in chick embryos using both morpholino-mediated knock-down as well as CRISPR/Cas9 knock-out using three different gRNAs. We then let the embryos develop for an additional one (for gene expression) or two (for staging and migration) days and analyzed several potentially affected biological processes. Surprisingly, we noticed that HIF-2α knockdown embryos were developmentally delayed compared with their control counterparts (**Figure 3A-D**). The stages of embryos following CRISPR/Cas9- or morpholino mediated loss of HIF-2α were determined by their Hamburger and Hamilton developmental stage *in ovo* (**Figure 3A-B**) and by counting somites *ex ovo* (**Figure 3C-D**).

**Fig. 3.**
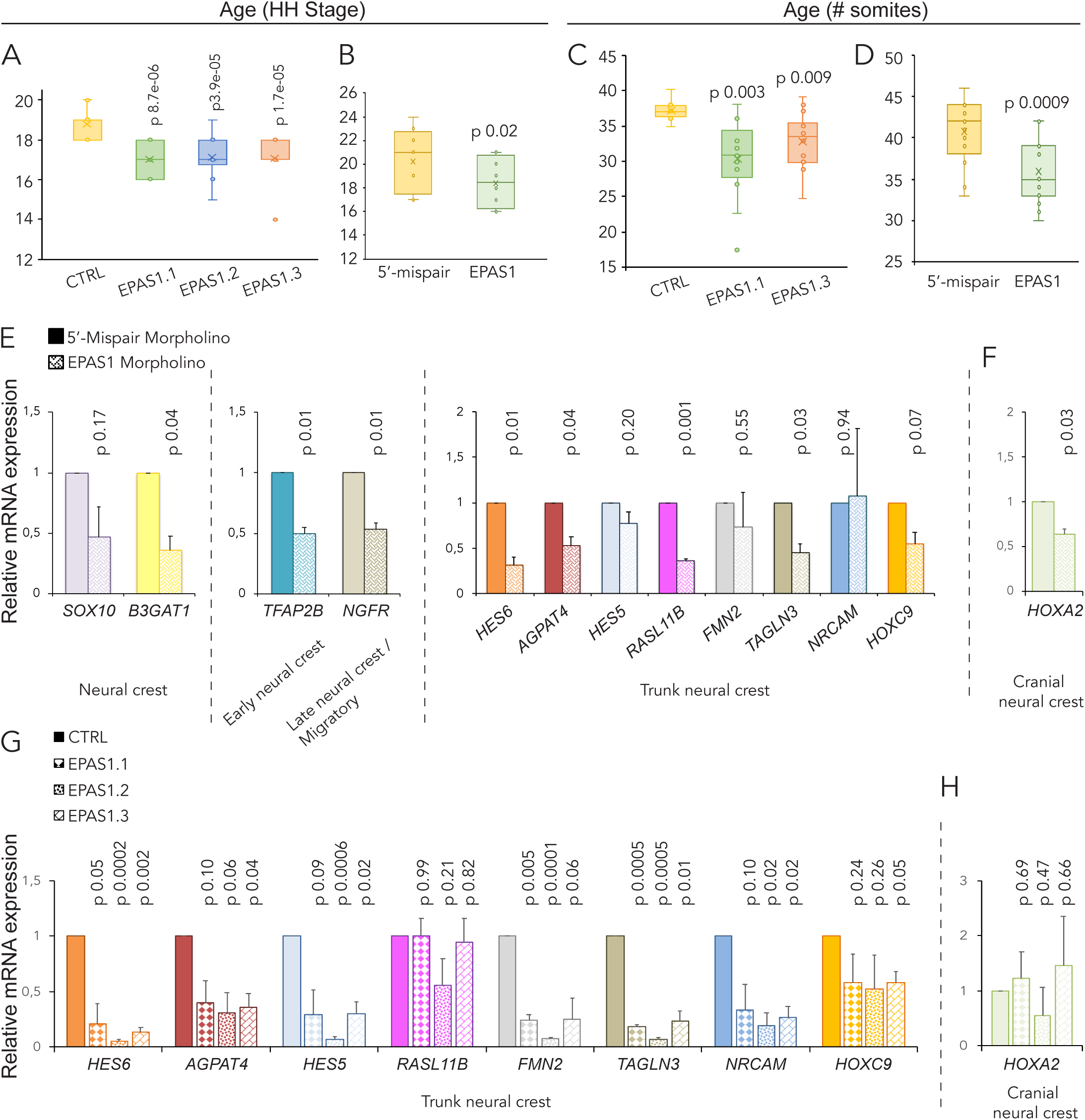
Knockdown of HIF-2α delays embryogenesis. **A.** Hamburger Hamilton (HH) staging of embryos 36 hours post-electroporation with a non-targeting (CTRL) gRNA compared to three different gRNAs targeting *EPAS1* (EPAS1.1, EPAS1.2, EPAS1.3) by head- and tail morphology. Number of embryos analyzed were n=14 (CTRL), n=10 (EPAS1.1), n=14 (EPAS1.2) and n=14 (EPAS1.3). Statistical significance was determined by one-way ANOVA comparing non-targeting CTRL with each individual *EPAS1* gRNA. **B.** Hamburger Hamilton (HH) staging of embryos 44 hours post-electroporation with 5’-mispair or *EPAS1* targeting morpholinos by head- and tail morphology. Number of embryos analyzed were n=20 (5’-mispair), n=16 (EPAS1). Statistical significance was determined by one-way ANOVA. **C.** Determination of embryonic age by number of somites 36 hours post-electroporation. Number of embryos analyzed were n=8 (CTRL), n=13 (EPAS1.1) and n=14 (EPAS1.3). Statistical significance was determined by one-way ANOVA comparing non-targeting CTRL with each individual *EPAS1* gRNA. **D.** Determination of embryonic age by number of somites 44 hours post-electroporation. Number of embryos analyzed were n=17 (5’-mispair), n=15 (EPAS1). Statistical significance was determined by one-way ANOVA. **E-F.** Relative mRNA expression of /trunk/ neural crest **(E)** and cranial neural crest **(F)** associated genes in trunk neural crest cells derived from embryos electroporated with 5’-mispair or *EPAS1* morpholinos, measured by qRT-PCR 24 hours post-electroporation. **G-H.** Relative mRNA expression of trunk neural crest **(G)** and cranial neural crest **(H)** associated genes in trunk neural crest cells derived from embryos electroporated with non-targeting CTRL or three *EPAS1* gRNAs, measured by qRT-PCR 24 hours post-electroporation. **E-H.** Data presented as mean of n=2 biologically independent repeats, error bars denote SEM. Statistical significance was determined by two-sided student’s t-test (**E-H**), comparing non-targeting CTRL with each individual *EPAS1* gRNA in **G-H**.

Electroporation efficiency was confirmed by analyzing *EGFP* expression (Supplementary Figure S2A-B). Knockdown of HIF-2α, either by morpholino or CRISPR/Cas9, led to decreased expression levels of genes representative of early and migrating neural crest as well as trunk neural crest cells in particular (Frith et al., 2018; Murko, Vieceli, & Bronner, 2018) (**Figure 3E-G**, respectively, and Supplementary Figure S2C-D). In contrast, the cranial neural crest associated gene *HOXA2* was not affected (**Figure 3F, H**).

One of the most important features of neural crest cells is their migratory ability. Trunk neural crest cells destined to form the sympathetic chain ganglia migrate ventrally. After HIF-2α loss of function using either morpholinos or CRISPR/Cas9, HNK1 positive migratory neural crest cells were detected on the control side in all embryos (**Figure 4A-D**) as well as on the side electroporated with non-targeting gRNA CTRL and control 5’-mismatch morpholino (**Figure 4A** **and** 4C, respectively). In contrast, loss of HIF-2α profoundly reduced the numbers of HNK1 positive cells migrating to ventral regions of the embryo (CRISPR/Cas9, **Figure 4B**; morpholino, **Figure 4D**).

**Fig. 4.**
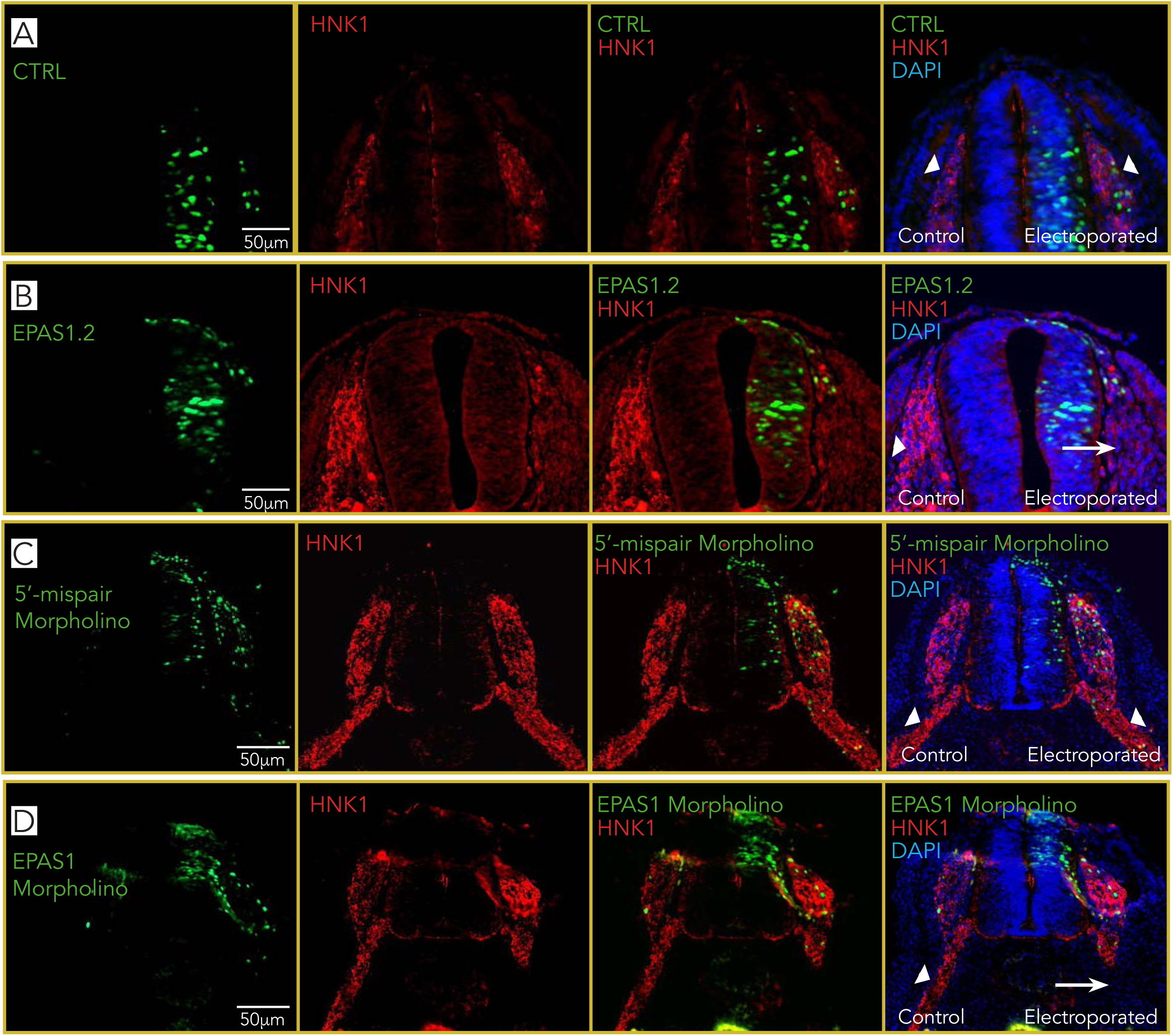
Knockdown of HIF-2α affects migration of trunk neural crest cells. **A-D.** Immunostaining of HNK1 (red) marking migrating crest cells in one-sided electroporated embryos (right side). Electroporated cells (non-targeting CTRL gRNA **(A)**, gRNA #2 targeting *EPAS1* (EPAS1.2; **B**), 5’-mispair morpholino **(C)** or *EPAS1* morpholino **(D)**) are seen in green. DAPI was used to counterstain nuclei. Embryo sections from trunk axial level are from 48 hours (**A-B**) or 44 hours (**C-D**) post-electroporation.

SOX9, a member of the SoxE family of transcription factors, is important for neural crest fate. It is expressed in premigratory neural crest cells at all axial levels and promotes their lineage progression. Importantly, transverse sections through the trunk of embryos electroporated with control or two different EPAS1 targeting gRNA constructs showed no differences in SOX9 expression (Supplementary Figure S3A-C), suggesting that neural crest lineage specification was unaffected by loss of HIF-2α.

### Over-expression of HIF-2α has similar effects as loss-of-function

Similar to the loss-of-function experiments, overexpression of HIF-2α led to delayed embryonic development (**Figure 5A**) and perturbed migration as visualized by HNK1 staining (**Figure 5B**). Expression of neural crest- and trunk specific genes was slightly suppressed (**Figure 5C** and Supplementary Figure S4A) whereas expression of cranial neural crest gene *HOXA2* was slightly induced (**Figure 5C**). Overexpression of *EPAS1* was confirmed by qRT-PCR (Supplementary Figure S4B).

**Fig. 5.**
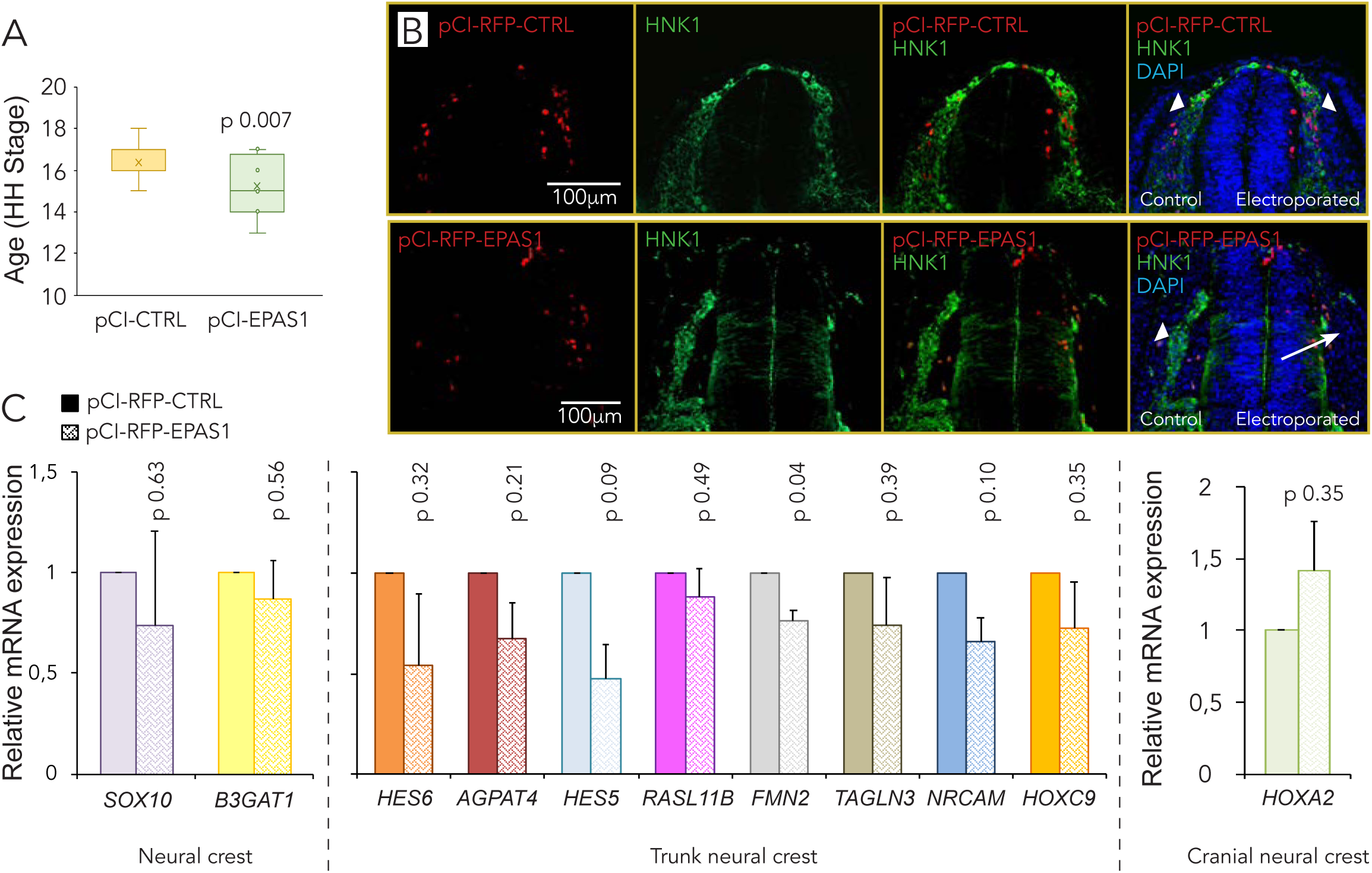
Controlled expression of HIF-2α is required to maintain embryonic homeostasis. **A.** Hamburger Hamilton (HH) staging of embryos 24 hours post-electroporation with a control (pCI-CTRL) or *EPAS1* overexpression construct (pCI-EPAS1), determined by head- and tail morphology. Number of embryos analyzed were n=16 (CTRL), n=20 (EPAS1). Statistical significance was determined by one-way ANOVA. **B.** Immunostaining of HNK1 (green) marking migrating crest cells in one-sided electroporated embryos (right side). Electroporated cells (CTRL or EPAS1) are seen in red. DAPI was used to counterstain nuclei. Embryo sections from trunk axial level are taken 48 hours post-electroporation. **C.** Relative mRNA expression of /trunk/ neural crest and cranial neural crest genes in trunk neural crest cells derived from embryos electroporated with CTRL or EPAS1 vectors, measured by qRT-PCR 24 hours post-electroporation. Data presented as mean of n=2 biologically independent repeats, error bars denote SEM. Statistical significance was determined by two-sided student’s t-test.

### Trunk neural crest cells proliferate extensively in response to dysregulated HIF-2α

We next examined cell proliferation in premigratory and migrating neural crest cells after loss of HIF-2α using real-time EdU pulse chase labeling optimized for avian embryos (Warren et al., 2009). Quantifying the proportion of premigratory and early migrating neural crest cells that had incorporated EdU demonstrated a significant induction of proliferating cells with an average proportion of double positive cells of 22% and 70% in the 5’-mismatch versus EPAS1 morpholino targeted embryos, respectively (p 0.029; **Figure 6A-B**).

**Fig. 6.**
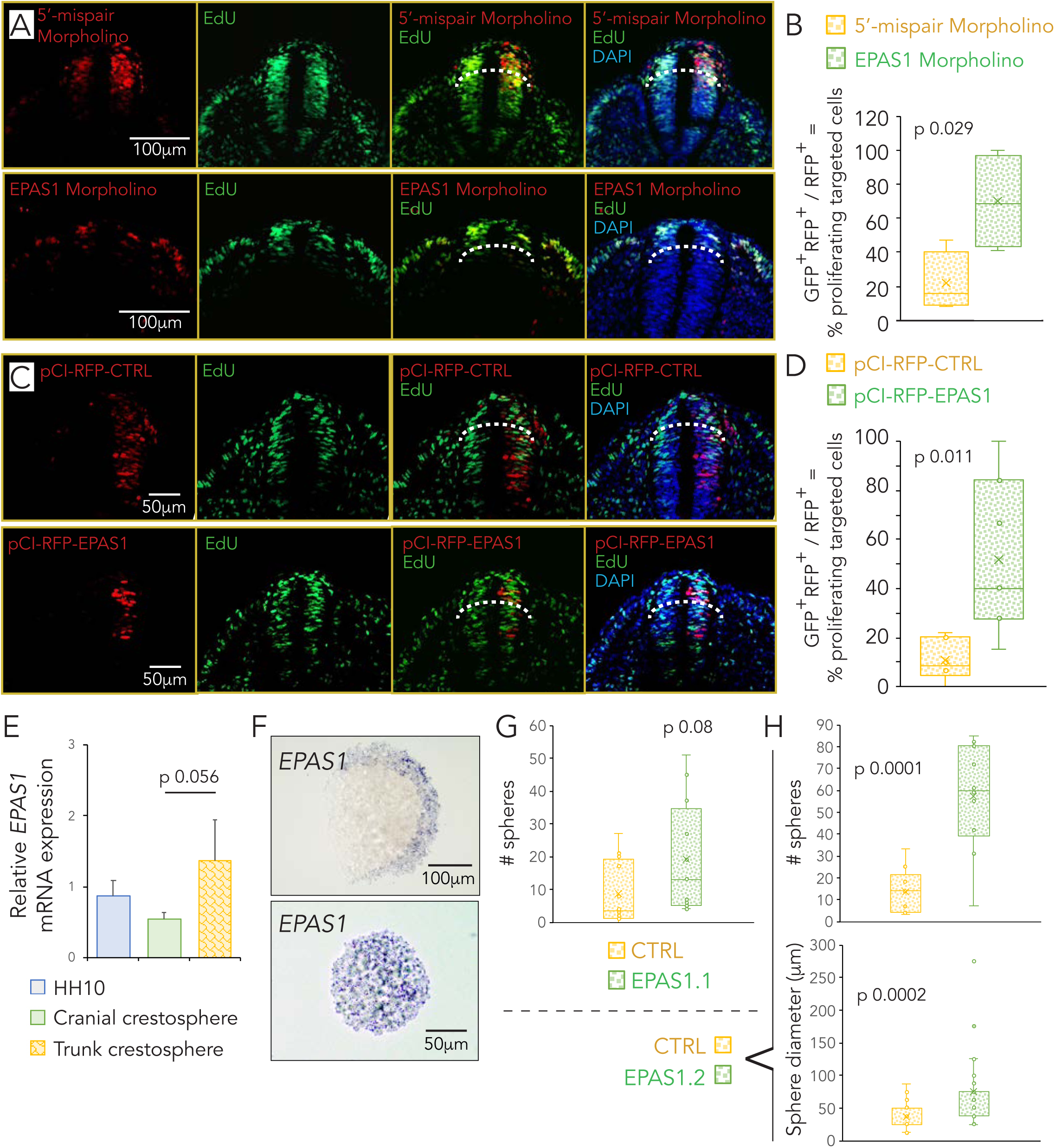
Knockdown of HIF-2α affects proliferation and stemness of trunk neural crest cells. **A-D.** Embryo sections from trunk axial level after real-time EdU labeling. Proliferating EdU^+^ cells are seen in green and electroporated cells (5’-mispair and *EPAS1* morpholinos (**A**); pCI-CTRL and pCI-EPAS1 (**C**)) are seen in red. DAPI was used to counterstain nuclei. Quantification of proliferating cells was performed by manual counting of RFP^+^ only as well as double positive cells. Only construct targeted trunk neural crest cells (above dotted line) were included. Number of cells analyzed were n=82 (5’-mispair morpholino) and n=303 (EPAS1 morpholino) **(B)**; n=211 (pCI-CTRL) and n=139 (pCI-EPAS1) **(D)**. **E.** Relative mRNA expression of *EPAS1* in wild type HH10 embryos (blue bar) and crestosphere cells established from the cranial axial level (green bar) or trunk axial level (yellow bar) measured by qRT-PCR. Expression is presented as mean of n=4 (cranial) or n=3 (trunk) biological replicates and error bars represent SEM. Expression difference between cranial and trunk crestospheres, p=0.056, as determined by two-sided student’s t test. Expression in wild type HH10 embryos is presented as mean of n=3 technical replicates. **F.** *In situ* detection of *EPAS1* mRNA in trunk derived crestospheres. **G.** Primary sphere assay, i.e. quantification of self-renewal from crestospheres established from dissociated trunk neural tubes of HH13+/14-embryos previously electroporated *in ovo* at HH10^+^/HH11 with non-targeting gRNA (CTRL) or gRNA targeting *EPAS1* (EPAS1.1). One cell/well (n=10 wells per group) were seeded at T= 0 days. Number of spheres were manually counted in each well after T= 1 week. Statistical significance was determined by one-way ANOVA. **H.** Quantification of self-renewal (as described in **G.**) and sphere size from crestospheres established from dissociated trunk neural tubes of HH13+/14-embryos electroporated with non-targeting gRNA (CTRL) or gRNA targeting *EPAS1* (EPAS1.2). Sphere size by manual measurements converted to factual unit (μm). Statistical significance was determined by one-way ANOVA.

After over-expression of HIF-2α, real-time EdU incorporation demonstrated that cells with increased expression of HIF-2α also became highly proliferative with an average proportion of double positive cells of 11% and 52% in the control and EPAS1 overexpressing embryos, respectively (p 0.011; **Figure 6C-D**). We conclude that neural crest proliferation is highly sensitive to expression levels of HIF-2α, suggesting that levels must be tightly controlled for proper development.

### HIF-2α downregulation enhances stem cell properties of trunk NC cells

Neural crest-derived crestosphere cultures (Mohlin & Kerosuo, 2019; Mohlin, Kunttas, et al., 2019) enable studies on stemness properties of these cells *in vitro*. Therefore, we examined *EPAS1* expression in crestosphere cultures, in which multipotent neural crest cells can be maintained in a stem cell-like state *in vitro* (Kerosuo et al., 2015; Mohlin, Kunttas, et al., 2019). When comparing crestosphere cultures derived from trunk versus cranial axial levels, we noted that *EPAS1* was enriched in trunk crestospheres (**Figure 6E**). *In situ* hybridization further revealed two separate patterns of *EPAS1* expression in trunk crestospheres: equal distribution throughout the spheres or concentration in cells at the edges of the spheres (**Figure 6F**).

Next, we established trunk crestospheres from embryos electroporated with a control gRNA construct or two different gRNAs targeting *EPAS1*. Primary sphere assays demonstrated that cells with dysregulated HIF-2α levels had an increased ability to form new spheres when seeded as single cells (1 cell/well; **Figure 6G-H**). In addition, crestosphere cultures derived from embryos electroporated with the *EPAS1* targeting construct formed larger spheres compared to their control counterparts (**Figure 6H**).

### RNA-seq after loss of HIF-2α in neural crest cells identifies downstream genes associated with invasion, growth arrest and developmental regulation

To investigate gene expression changes after loss of HIF-2α, we performed loss of function experiments at premigratory stages of neural crest development (HH10^+^/HH11 in avian embryos) using a splice targeting morpholino as above. Neural tubes from trunk region were dissected 24 hours post-electroporation (at stage ∼HH16) and subsequently analyzed by RNA sequencing. Correlation plot of all genes from the dataset demonstrated that trunk neural crest cells after knockdown of HIF-2α indeed differ from those injected with control scrambled morpholino (spearman p>0.96; **Figure 7A**). Setting a cut-off at p<0.005 and removing all hits that were not annotated (NA), we identified 97 genes of interest (**Figure 7B**). The top ten genes down- and upregulated (assessed by log2 fold differences in expression) by knockdown of HIF-2α are summarized in **Figure 7C**, while the complete list of these 97 genes can be found in Supplementary Table S1.

**Fig. 7.**
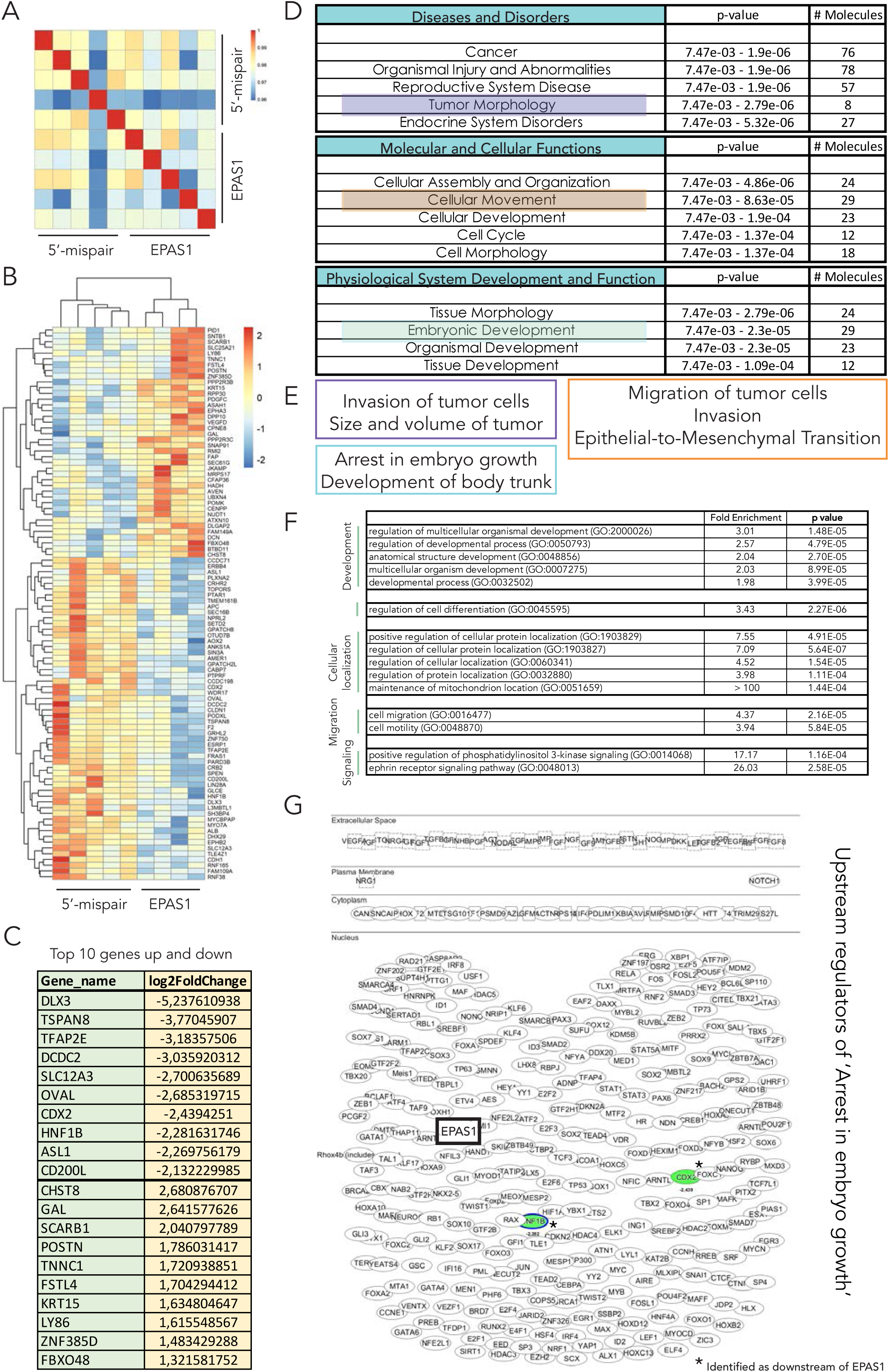
Gene set enrichment analysis identifies HIF-2α downstream affected processes. **A-B.** Hierarchical clustering of significantly Differentially Expressed Genes (DEGs;cut-off p<0.005) identified from RNA sequencing comparing 5’-mispair and *EPAS1* morpholino samples. **C.** List of the top ten upregulated and top ten downregulated genes from the RNA sequencing data. **D.** Hypothesis-free/exploratory analysis of the 97 DEGs using IPA (Fishers Exact Test for the range of p-value calculation) revealed a series of top five hits (p<0.05) in the respective categories “Disease and Disorders”, “Molecular and Cellular Functions” and “Physiological System Development and Function” downstream processes**. E.** Deeper analysis of processes identified in **(D)**. **F.** Selected list of enriched cellular processes from Panther analyses. Complete list can be found in **Supplemental Table S2**. **G.** Deeper analysis of potential upstream regulators of the “arrest in embryo growth” process identified in **E**. The shape of molecules and their meaning, i.e. correspondence to protein family etc., is found here: http://qiagen.force.com/KnowledgeBase/KnowledgeIPAPage?id=kA41i000000L5rTCAS. As an example, the diamond shaped molecules correspond to enzymes, oval standing shapes should be read as transmembrane receptors and lying oval shapes are transcription regulators. Green nodes indicate down-regulated molecules. The intensity of the color reveals the strength of the expression i.e. the stronger the color the more significant.

Gene set enrichment analysis (GSEA) on the RNA sequencing data described above demonstrated that two out of the top five processes connected to disease were cancer and tumor morphology (with 29 and 8 out of 97 molecules, respectively; **Figure 7D**). Deeper analysis of tumor morphology showed that genes associated with invasion of tumor cells and size and volume of tumor were particularly enriched, i.e. these associated genes linked to specific disease categories are not due to random chance but are statistically significant (p<0.05) (**Figure 7E**). Consistent with *in vivo* data, we identified cellular movement as one of the top molecular and cellular functions affected, with invasion as well as migration of tumor cells and epithelial-to-mesenchymal transition as predicted downstream pathways (**Figure 7E**). GSEA also revealed enrichment of genes associated with arrest in embryo growth (**Figure 7D-E**). We conclude that the predicted cellular functions derived from our RNA sequencing experiment overlap with *in vivo* data (cf. **Figures 3**-**6**). Top networks from the RNA sequencing data showed enrichment of two signaling pathways, the ephrin receptor- and phosphatidylinositol 3-kinase (PI3K) signaling pathways (**Figure 7F** and Supplementary Figure S5A, with full list of gene ontology enriched processes in Supplementary Table S2).

Dividing the hits from RNA sequencing data that overlap with genes enriched for migration of tumor cells revealed a large subset of genes that encode for plasma membrane associated- or are secreted proteins (Supplementary Figure S5B). Several of these overlapping genes were among the 97 significantly differentially expressed (with cut-off p<0.005), suggesting a close regulatory relationship between HIF-2α and migration at least during these time points of development.

Given the effects we observed on embryonic development *in vivo*, we mapped potential upstream regulators of arrest in embryo growth. As expected, most genes were transcription factors, including *EPAS1* itself (**Figure 7G**). Among the predicted upstream regulators of arrested growth, genes associated with stem cells, BMP signaling and EMT were highly enriched (**Table 1** and Supplementary Table S3).

**Table 1.**
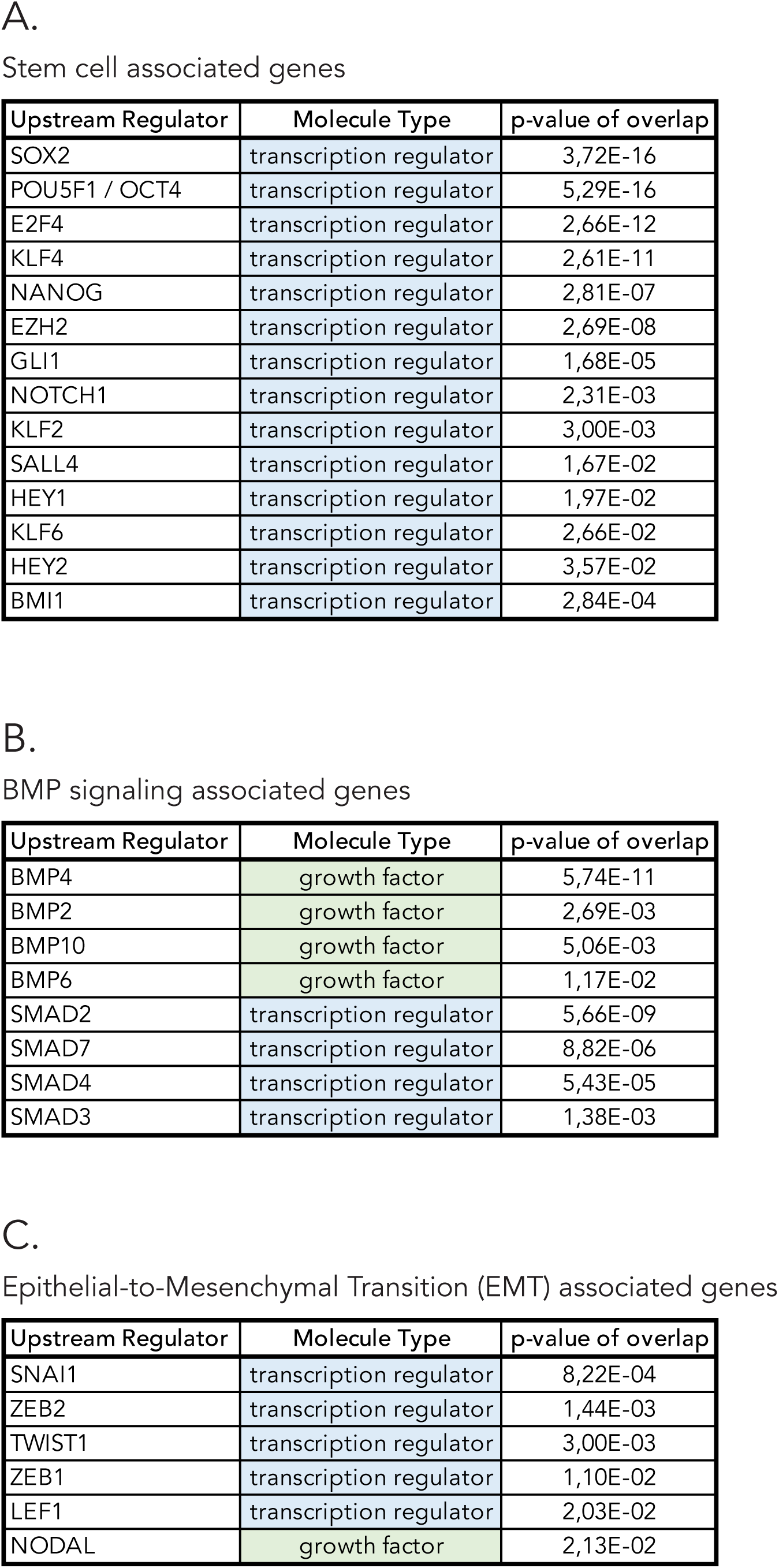
Selected genes identified as potential upstream regulators of arrested embryo growth. Genes associated with stem cells **(A)**, BMP signaling **(B)** and EMT **(C)** were particularly enriched.

Two other predicted genes upstream of arrested embryo growth were *CDX2* and *HNF1B*, also among the 97 significantly (cut-off p<0.005) differentially expressed in the RNA sequencing data. Deeper analysis of these genes revealed autocrine signaling as well as an interconnected regulation between the two (Supplementary Figure S5C). EMT related genes *ZEB2* and *SNAI1* are negatively regulated by both of these genes (Supplementary Figure S5C). In addition, *CDX2* was predicted to regulate *MYCN*, a transcription factor commonly amplified in aggressive neuroblastoma (Supplementary Figure S5C). Of the significantly (cut-off p<0.005) differentially expressed genes, *CDX2* and *HNF1B* were predicted to be upstream regulators of *EPAS1*. The majority of predicted *EPAS1* upstream regulators were transcription factors, and we observed an enrichment for stem cell associated genes (Supplementary Table S4).

### Trunk neural crest associated genes are enriched in neuroblastoma

Neuroblastoma has long been recognized as derived from sympathetic neuroblasts of trunk neural crest based on marker expression and tumor localization (De Preter et al., 2006; Hoehner et al., 1996). However, recent studies from Adameyko and colleagues (Furlan et al., 2017; Kastriti et al., 2019; Soldatov et al., 2019) have raised questions regarding the origin of chromaffin cells as well as neuroblasts during embryonic development. While chromaffin cells mainly derive from Schwann cell precursors (Furlan et al., 2017), sympathetic neuroblasts are derived from sympathoadrenal precursor cells (Kastriti et al., 2019). Using a recently published dataset of migratory trunk neural crest enriched genes (Murko et al., 2018) as well as established neural crest and developmental markers, we examined connections between neuroblastoma and trunk neural crest cells. We compared expression of early neural crest marker *TFAP2B* as well as trunk neural crest markers *RASL11B*, *TAGLN3*, *NRCAM*, *AGPAT4*, *FMN2*, *HES5*, *HES6* (Murko et al., 2018) and *HOXC9* (Frith et al., 2018) in cancer cell lines of different origins (Cancer Cell Line Encyclopedia (CCLE) containing >600 cell lines; cancer types with n*≥*4 cell lines were selected for further analysis (R2; http://hgserver1.amc.nl)) demonstrating enriched expression for the majority of these genes in neuroblastoma cells as compared to other cancer types (**Figure 8A** and Supplementary Figure S6A-C). Cranial neural crest marker *HOXA2* was on the other hand not enriched in neuroblastoma as compared to other cancer types (Supplementary Figure S6D). Neuroblastoma patient-derived xenograft (PDX) cells have been established from mouse models of orthotopic implantation of patient-derived tumor pieces (Braekeveldt et al., 2015; Persson et al., 2017). These PDX cells retain characteristics of their respective patient tumor and metastasize to clinically relevant sites *in vivo*. Real-time quantitative PCR analyses demonstrated significant enrichment of neural crest (*TFAP2B*, *SOX10*) and trunk neural crest (*RASL11B*, *FMN2*, *TAGLN3*, *NRCAM*, *HES6*, *HES5*, *AGPAT4*) gene expression in neuroblastoma PDX cells as compared to cells from renal cell carcinoma (RCC-4 and 786-0) and liver cancer cell lines (Hep3b) (**Figure 8B-C** and Supplementary Figure S6E).

**Fig. 8.**
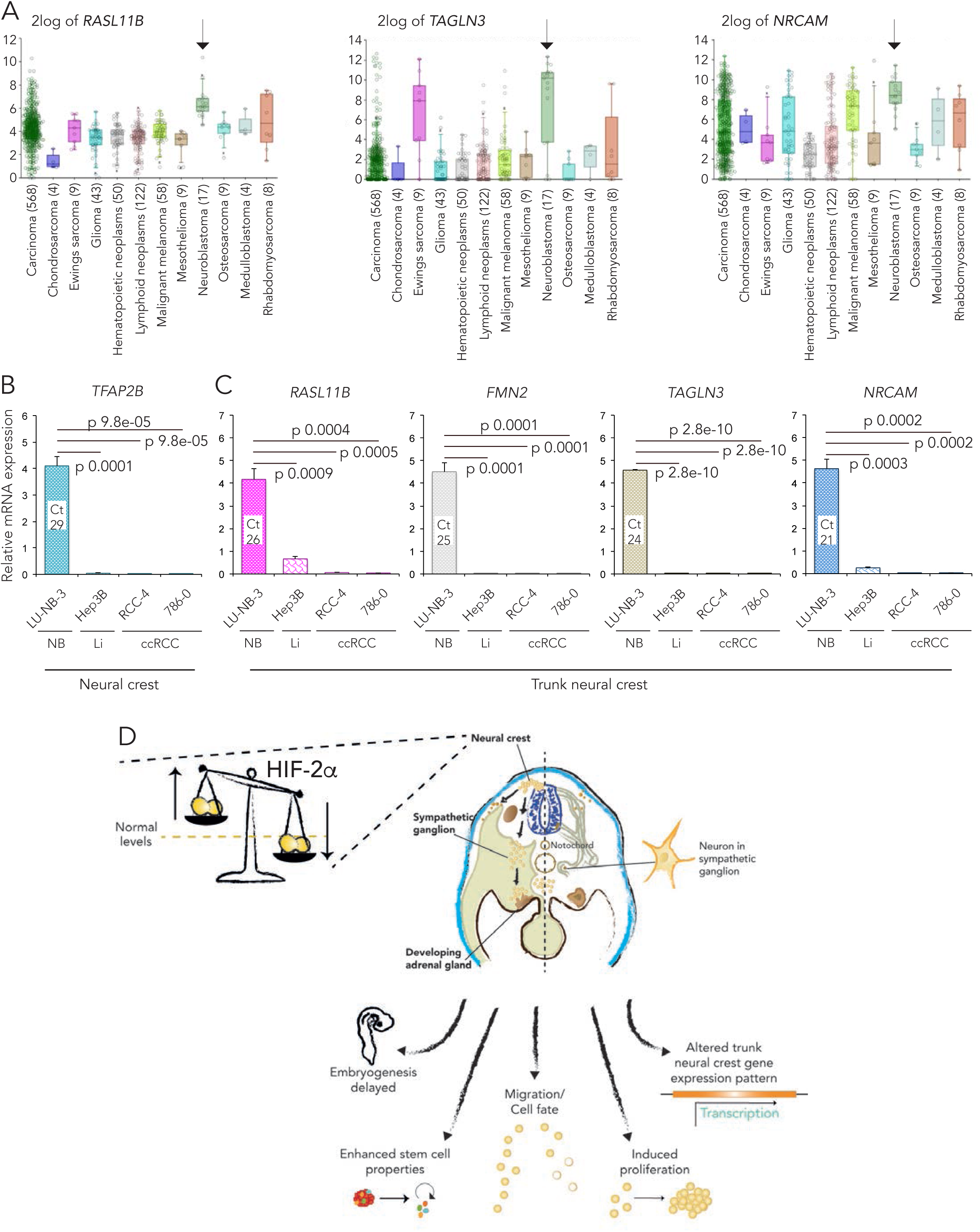
Trunk neural crest associated genes are enriched in neuroblastoma. **A.** Trunk neural crest (*RASL11B*, *TAGLN3* and *NRCAM*) gene expression in cancer types of different tissue origins. Data from the Cancer Cell Line Encyclopedia (CCLE) dataset, tissue origin with samples n>3 were chosen for further analysis. Number in brackets represents the number of cell lines from each tissue origin. Arrows highlight neuroblastoma. **B-C.** Relative mRNA expression of neural crest (*TFAP2B*, **(B)**) and trunk neural crest (*RASL11B*, *FMN2*, *TAGLN3* and *NRCAM* **(C)**) genes measured by qRT-PCR. Expression in LU-NB-3 neuroblastoma (NB) patient-derived xenograft cells were compared to liver cancer (Li) Hep3B and clear cell renal cell carcinoma (ccRCC) RCC-4 and 786-0 cell lines. Data are presented as mean of n=3 biologically independent replicates and error bars represent SEM. Statistical significance comparing Hep3B, RCC-4 or 786-0 individually to LU-NB-3 was tested using two-sided students *t test*. **D.** Schematic summary of developmental effects following dysregulated HIF-2α expression levels in trunk neural crest cells.

## Discussion

It has long been assumed that the childhood tumor form of neuroblastoma derives from sympathoadrenal neuroblasts. These assumptions have been based on the expression of proteins in neuroblastoma that are also expressed by embryonic sympathetic neurons during normal development, as well as the location where these tumors arise (i.e. along the sympathetic ganglia). HIF-2α has been implicated in tumor growth and is expressed in cancer stem cells of several tumors including neuroblastoma. However, little has been known about its expression and function during normal development. Here, we show that the HIF-2α protein is expressed in trunk neural crest cells and sympathetic neuroblasts during normal embryogenesis in three different species: human, mouse and avian and examine its function using the chick embryo as a model amenable to experimental manipulation. Comparable data across human, mouse and avian tissue suggest that cross-species interpretation of further results is valid.

Either knock-down or overexpression of HIF-2α in premigratory chick trunk neural crest affects several important functions. Not only do embryos with dysregulated HIF-2α have developmental delays compared to controls, but they also exhibit altered neural crest gene expression profiles. Consistent with observed *in vivo* effects, RNA sequencing demonstrates a global genome level change after loss of HIF-2α, with upregulation of genes involved in invasive behavior and growth arrest. Furthermore, we observed altered trunk neural crest migratory patterns as well as enhanced proliferative capacity of trunk neural crest cells *in vivo*, as well as in our RNA sequencing data.

Despite extensive proliferation of trunk neural crest cells with dysregulated HIF-2α expression, the embryos as a whole develop at a slower pace than their control counterparts. In general, cell division of trunk neural crest cells is limited during their active migratory phase. We speculate that the observed embryonic delays relative to increased trunk neural crest cell proliferation may be the result of a skewed cell division to migration ratio, with increased proliferation perhaps causing a failure in cell migration.

The capacity to self-renew is an important feature of stem-like cells. Our data suggest that *EPAS1* knockout cells exhibit enhanced self-renewal, in line with observations in neuroblastoma cells with aberrant HIF-2α expression which are more immature and neural crest-like (Pietras et al., 2008). In addition, crestospheres formed by HIF-2α dysregulated single cells were larger, a sign of enhanced proliferative capacity in agreement with our EdU results.

The RNA sequencing data revealed enrichment of two signaling pathways, the ephrin receptor- and PI3K pathways. This suggests that environmental cues may be influencing trunk neural crest behavior. Of note, we have recently identified that PI3K-mTORC2 regulates HIF-2α expression and functions as a valid treatment target in neuroblastoma (Mohlin et al., 2015; Mohlin, Hansson, et al., 2019). Genes associated with migration of tumor cells mainly encode for plasma membrane and secreted proteins, including several members of the matrix metalloproteinase (MMP) family. MMPs promote invasion and migration by degrading components of the extracellular matrix and have been shown to be regulated by HIF-2α in several different tumor forms (Koh, Lemos, Liu, & Powis, 2011; Petrella, Lohi, & Brinckerhoff, 2005), further reinforcing a possible connection between HIF-2α, trunk neural crest cells and invasive behavior.

The stem cell gene *POU5F1*, more commonly known as Oct4, is driven by HIF-2α in immature cells during development (Covello et al., 2006). We found that Oct4 is predicted to be upstream of arrested embryo growth, but also an upstream regulator of *EPAS1* itself. One of the *EPAS1* target molecules connecting Oct4 and HIF-2α is *CDX2*, which in turn is upstream of *EPAS1* as well as arrested embryo growth (**Supplementary Figure S5C**). CDX2 is indeed one of the major players involved in mediating the HIF-2α driven effects on embryonic development and considering that CDX2 is an early trunk neural crest marker (Frith et al., 2018), a possible explanation for delayed embryonic development might be halted trunk neural crest commitment.

These findings contribute to understanding of a complex regulatory network involved in mediating trunk neural crest development. We posit that the cancer associated protein HIF-2α may play a central role in embryonic growth, global gene expression, migration, proliferation and stem cell features of neural crest cells within this network (**Figure 8D**). Moreover, our results highlight the importance of careful regulation of HIF-2α levels for maintenance of normal embryonic growth and differentiation.

## Materials and Methods

### Chick embryo tissue

According to Swedish regulations (Jordbruksverkets föreskrift L150, §5) work on chick embryos younger than embryonic day 13 do not require Institutional Animal Care and Use Committee oversight.

### Human and mouse fetal tissue

Human fetal tissue (ethical approval Dnr 6.1.8-2887/2017, Lund University, Sweden) was obtained from elective abortions. Tissue samples were dissected in custom-made hibernation medium (Life Technologies) and fixed in 4% formaldehyde overnight. Following a sucrose gradient, embryos were embedded in gelatin for transverse sectioning at 12μm (ew5) or 7μm (ew6) using a cryostat.

### Embryos and perturbations

Chick embryos were acquired from commercially purchased fertilized eggs and incubated at 37.5°C until desired developmental Hamburger Hamilton (HH) stages were reached (Hamburger & Hamilton, 1951). Optimal conditions for high transfection efficiency applying one-sided electroporation *in ovo* were determined to 5 pulses of 30ms each at 22V. Ringer’s balanced salt solution (Solution-1: 144g NaCl, 4.5g CaCl•2H_2_O, 7.4g KCl, ddH_2_O to 500ml; Solution-2: 4.35g Na_2_HPO_4_•7H_2_O, 0.4g KH_2_PO_4_, ddH_2_O to 500ml (adjust final pH to 7.4)) containing 1% penicillin/streptomycin was used in all experiments. Morpholinos used were from GeneTools with the following sequences; splice targeting EPAS1 oligo (5’-GAAAGTGTGAGGGAACAAGTTACCT-3’) and a corresponding 5’-mispair oligo (5’-GAtAcTGTcAGGcAACAAcTTACCT-3’). Morpholinos were injected at a concentration of 1mM and co-electroporated with a GFP tagged empty control vector (1 ug/ul). RFP-tagged *EPAS1* overexpression construct and corresponding empty control vector were electroporated at a concentration of 2.5 ug/ul. CRISPR constructs with gRNA non-targeting control (#99140, Addgene) or gRNAs targeting *EPAS1* (EPAS1.1.gRNA Top oligo – 5’ ggatgGCTCAGAACTGCTCctacc 3’, Bot oligo – 5’ aaacggtagGAGCAGTTCTGAGCc 3’; EPAS1.2.gRNA Top oligo – 5’ ggatgAAGGCATCCATAATGCGCC 3’, Bot oligo – 5’ aaacGGCGCATTATGGATGCCTTc; 3’; EPAS1.3.gRNA Top oligo – 5’ ggatgAAATACATGGGTCTCACCC 3’, Bot oligo – 5’ aaacGGGTGAGACCCATGTATTTc3’) were cloned into U6.3>gRNA.f+e (#99139, Addgene) and electroporated at a concentration of 1.5 ug/ul, and accompanying Cas9 (#99138, Addgene) at 2 ug/ul (Gandhi, Haeussler, Razy-Krajka, Christiaen, & Stolfi, 2017). Embryos were allowed to sit at room temperature for 8 – 10 hours in order to allow the Cas9 protein to fold before further incubation of the embryos at 37.5°C.

For harvesting of trunk neural crest cells for RNA extraction, embryos were incubated at 37.5°C for 24 (morpholinos and overexpression vectors) or 36 (CRISPR/Cas9) hours post-electroporation. Embryos were incubated for 24 to 48 hours post-electroporation before dissecting whole embryos for fixation and embedding.

### Cloning

To overexpress HIF-2α, the gallus gallus *EPAS1* coding sequence was amplified using the following primers; Fwd: 5’AAACTCGAGGCCACCATGGACTACAAAGACGATGACGACAAGGCAGGTATGACAGCTGACAAGGAGAAG-3’, Rev 5’-AAAGCTAGCTCAGGTTGCCTGGTCCAG-3’ and cloned into the pCI H2B-RFP vector (Addgene plasmid #92398). For CRISPR/Cas9 targeting, oligos designed to target *EPAS1* at three different locations (EPAS1.1, EPAS1.2 and EPAS1.3) were annealed pairwise at a concentration of 100 μM per oligo using T4 DNA Ligase Buffer in dH_2_O by heating to 95°C for 5 minutes. The annealed oligo reactions were cooled to room temperature and diluted. The U6.3>gRNA.f+e (#99139, Addgene) vector was digested over night with BsaI-HF enzyme (New England Biolabs) and gel extracted. gRNAs were cloned into the digested U6.3>gRNA.f+e vector using T4 DNA Ligase (New England Biolabs) at room temperature for 20 minutes. Successful inserts were identified by colony PCR using U6 sequencing primer and gRNA reverse oligo specific to each *EPAS1* gRNA.

### Cell culture

The human neuroblastoma cell line SK-N-BE(2)c (ATCC; Manassas, VA, US) and hepatocellular carcinoma cell line Hep3b (ATCC; Manassas, VA, US) were cultured in MEM while renal cell carcinoma RCC4 and 786-O cell lines were cultured in DMEM, supplemented with 10% fetal bovine serum and 100 units penicillin and 10μg/mL streptomycin. As part of our laboratory routines, all cells were maintained in culture for no more than 30 continuous passages and regularly screened for mycoplasma. SK-N-BE(2)c cells were authenticated by SNP profiling (Multiplexion, Germany).

### Neural tube dissection

Neural tubes from respective axial levels were carefully dissected out from embryos at designated somite stages. For cranial-derived cultures, the very anterior tip was excluded, and the neural tube was dissected until the first somite level as previously described (Kerosuo, Nie, Bajpai, & Bronner, 2015). For trunk-derived cultures, the neural tube was dissected between somite 10-15 as previously described (Mohlin & Kerosuo, 2019; Mohlin, Kunttas, et al., 2019). Pools of neural tubes from 4 - 6 embryos were used for each culture.

### Crestosphere cell culture

Neural tube derived cells were cultured in NC medium (DMEM with 4.5g/L glucose (Corning), 7.5% chick embryo extract (MP Biomedicals; Santa Ana, CA, USA), 1X B27 (Life Technologies; Carlsbad, CA, US), basic fibroblast growth factor (bFGF, 20 ng/ml) (Peprotech; Stockholm, Sweden), insulin growth factor -I (IGF-I, 20 ng/ml) (Sigma Aldrich; Darmstadt, Germany), retinoic acid (RA; 60nM for cranial and 180nM for trunk, respectively) (Sigma Aldrich; Darmstadt, Germany), and 25 ng/ml BMP-4 (for trunk) (Peprotech; Stockholm, Sweden)) in low-adherence T25 tissue culture flasks as described previously (Mohlin & Kerosuo, 2019; Mohlin, Kunttas, et al., 2019).

### Self-renewal assay

Chick embryos at developmental HH stage 10+ were injected and electroporated with CRISPR/Cas9 constructs and allowed to develop at 37.5°C to reach HH stage 13/14. Crestosphere cultures were established from embryos electroporated with control, EPAS1.1 or EPAS1.2 constructs, respectively. Crestospheres were dissociated into single cells using Accutase (Sigma Aldrich; incubation at 37 °C for 40 min with one minute of pipetting every 10 min), and individual cells were manually picked using a p10 pipette tip under the microscope. Single cells were transferred to 96-well plates prepared with 100 μl of NC medium supplemented with retinoic acid and BMP-4 (Mohlin, Kunttas, et al., 2019). The absolute number of spheres formed in each well was quantified manually under the microscope. Five wells were analyzed per crestosphere culture. Sphere diameter was manually measured using the ImageJ software (spheres measured n=33 and n=27 for CTRL and EPAS1.2, respectively).

### EdU pulse chase labelling

Proliferation was measured using the Click-iT^TM^ EdU Cell Proliferation kit (Invitrogen #C10337) according to the manufacturer’s recommendations with optimizations from Warren et al (Warren, Puskarczyk, & Chapman, 2009). Chick embryos at developmental HH stage 10+ were injected and electroporated with morpholino or overexpression constructs and allowed to develop for additional 24 hours at 37.5°C. Eggs were then re-opened and EdU solution (500μM in PBS-DEPC) was added. Eggs were re-sealed and incubated at 37.5°C for another 4 hours before embryos were dissected in Ringer’s solution and fixed in 4% paraformaldehyde overnight. Embryos were washed in PBS-DEPC, H_2_O and 3% BSA in PBS-DEPC before permeabilization in 0.5% Triton-X. Embryos were hybridized in reaction cocktail (Click-iT Reaction buffer, CuSO_4_, Alexa Fluor 488 Azide and reaction buffer additive), washed and then DAPI stained. Embryos were after another round of washing processed through a sucrose gradient and embedded in gelatin.

### RNA sequencing

Chick embryos of stage HH10+ were injected with EPAS1 targeting or corresponding 5’-mispair morpholinos in the lumen of the neural tube and subsequently electroporated for construct uptake. Following 24 hours of incubation at 37.5°C, embryos were removed from the eggs in Ringer’s solution. Neural tubes from the trunk axial level of individual embryos were carefully dissected, removing surrounding mesodermal tissue, and transferred to Eppendorf tubes (neural tube tissue from one embryo per Eppendorf) that were snap frozen. RNA was extracted from each neural tube (5 samples per condition (EPAS1 and 5’-mispair, respectively)) using the RNAqueous Micro Kit (Ambion, #AM1931). Sequencing was performed using NextSeq 500 (Illumina). Alignment of reads was performed using the HISAT2 software and the reference genome was from the Ensemble database (Gallus gallus 5.0). Expression counts were performed using the StringTie software and differentially expressed genes (DEG) analysis was performed using DESeq2. To obtain a relevant working list out of the 1105 significantly DEGs, we set a cut-off at p<0.005 and removed all hits that were not annotated (NA), ending up with 97 genes. Significance (p values) were DESeq2 derived (Love, Huber, & Anders, 2014). RNA sequencing data have been deposited in NCBI’s Gene Expression Omnibus(Edgar, Domrachev, & Lash, 2002) and are accessible through GEO Series accession number GSE140319.

### Bioinformatics

Gene Set Enrichment Analysis (GSEA) for gene ontology, network and functional analyses were generated through the use of Panther database (analyses performed autumn 2018; (http://pantherdb.org/) (Thomas et al., 2003) together with the Ingenuity Pathway Analysis (IPA) software(Kramer, Green, Pollard, & Tugendreich, 2014) (QIAGEN Inc., https://www.qiagenbioinformatics.com/products/ingenuitypathway-analysis). The use of the two databases/software contributed to an added biological value in terms of knowledge. For a hypothesis-free/exploratory analysis of the 97 DEGs, IPA was used (p-value calculations using right-tailed Fisher Exact Test). However, IPA was mainly used for deeper exploration of the data where the biological hypotheses generated for the project were further explored. Here, a hypotheses-driven approach was taken where the following categories found from the IPA analysis of the 97 DEGs were further investigated; “Cellular Movement”, within the “Molecular and Cellular Function” result category, “Embryonic Development”, within the category “Physiological System Development and Function”, and “Tumor Morphology”, within the “Disease and Disorders” category. These three biological networks were further investigated within the data set at hand. The investigation for the possible overlap and connections between these networks in the context of the data were hence explored.

### Whole mount in situ hybridization of crestospheres

For whole mount *in situ* hybridization, crestospheres were fixed in 4% PFA for 30 minutes at RT and washed in DEPC-PBT. Samples were gradually dehydrated by bringing them to 100% MeOH and kept at -20°C until use. *In situ* hybridization was performed as previously described (Acloque, Wilkinson, & Nieto, 2008). Crestospheres were rehydrated back to 100% PBT, treated with Proteinase K/PBT, washed in 2 mg/ml glycine/PBT and post-fixed in 4% paraformaldehyde / 0.2% glutaraldehyde for 20 minutes. Crestospheres were then prehybridized in hybridization buffer for 2 hours at 70°C and hybridized with Digoxigenin (DIG)-labeled EPAS1 probe overnight at 70°C. Crestospheres were washed in Wash solution I and II (50% formamide, 1% SDS [Sodium Dodecyl Sulfate] and 5X SSC [NaCl and Na citrate] or 2X SSC, respectively), and blocked in 10% Sheep Serum for 2 hours followed by incubation with an anti-DIG antibody (1:2000) (Roche) in TBST / 1% sheep serum overnight at 4°C. On day 3, embryos were washed in TBST throughout the day and overnight. Crestospheres were washed in Alkaline phosphatase buffer (NTMT; 100mM NaCl, 100mM Tris-Cl (pH 9.5), 50mM MgCl2, 1%Tween-20) before visualizing the signal using 1-Step^TM^ NBT/BCIP Substrate Solution (ThermoFisher #34042). Stained crestospheres were fixed in 4% PFA for 20 minutes when they reached the desired state and dehydrated in MeOH to be stored at -20^°^C. Embryos were embedded in blocks of gelatin for transverse sectioning at 20 μm using a cryostat. Hybridization probe for avian *EPAS1* was prepared by using the following primers (Forward 5’-CAAGGAGAAGAAGAGGAGCA -3’; Reverse 5’-AAAGTGTGAGGAGGGCAAG -3’) and chick embryo cDNA as template. The amplified sequence was cloned into a pGEM-T Easy Vector before digestion and DIG RNA labeling (Roche #11277073910).

### Cryosections

Fixed embryos and crestospheres were incubated in a sucrose gradient (5% sucrose for 10 minutes and 15% sucrose for 10 minutes up to several hours) followed by incubation in 7.5% gelatin over night at 37°C. Embedded samples were cryosectioned at 7, 10, 12 or 20 μm.

### Immunohistochemistry and immunofluorescence

Immunohistochemistry on human (antigen retrieval by Target Retrieval Solution pH6.0 (DAKO #S1699)) and mouse fetal tissue was performed using Autostainer (Dako) and sections were counterstained with hematoxylin. Detection of HIF-2α by immunofluorescence was performed by incubation of embryo sections in ice cold acetone followed by 0.3% Triton-X in PBS. After washing in PBS, slides were blocked in DAKO serum-free ready-to-use block (DAKO, #X0909) for 1 hour before incubation with primary antibody (in DAKO antibody diluent with background reducing components (DAKO, #S3022)) overnight. Slides were washed in PBS and incubated with rabbit linker (DAKO, #K8019) followed by secondary antibody in 1% BSA/PBS. Detection of HNK1 and SOX9 by immunofluorescence was performed by blocking (10% goat serum and 0.3% Triton-X in TBST) of embryo sections followed by incubation with primary antibodies over night at +4°C. Slides were washed and incubated with secondary antibodies and DAPI for nuclear staining for 1 hour at RT before washing and mounting. Fluorescent images were acquired using an Olympus BX63 microscope, DP80 camera, and cellSens Dimension v 1.12 software (Olympus Cooperation). Detailed information on antibodies can be found in Supplemental Table S5.

### RNA extraction and quantitative real-time PCR

Total RNA was extracted using the RNAqueous Micro Kit (Ambion, #AM1931). Wild type whole embryos were carefully mechanically dissociated before lysis, pooling 2 to 4 embryos for each developmental stage. cDNA synthesis using random primers and qRT-PCR was performed as previously described (Mohlin et al., 2015). Relative mRNA levels were normalized to expression of two (avian; *18S, 28S*) or three (human; *UBC*, *SDHA*, *YWHAZ*) reference genes using the comparative Ct method (Vandesompele et al., 2002). Detailed information of primer sequences can be found in Supplementary Table S6.

### Fractionation and western blot

Cytoplasmic and nuclear extraction of proteins was performed using the NE-PER Nuclear and Cytoplasmic Extraction Reagents (Thermo Scientific). Proteins were separated by SDS-PAGE and transferred to HyBond-C-Extra nitrocellulose membranes. Detailed information on antibodies can be found in Supplemental Table S5.

### Oxygen sensing

Oxygen concentrations were measured through the trunk region of developing chick embryos *ex ovo* using microsensors in a flow system of MQ water. Microprofiles were measured in 50 embryos in developmental stages HH10 to HH24. Embryos were removed from the egg using filter paper as described in Mohlin and Kerosuo (Mohlin & Kerosuo, 2019), submerged in a plate with constant flow of newly shaken MQ of room temperature, and immediately measured. Oxygen microsensors were constructed and calibrated as described by Revsbech and Andersen (Revsbech & Andersen, 1989), mounted on a micromanipulator. The microsensor was manually probing the trunk region and data logged every second. Within the microprofile, ten consecutive data points of the lowest oxygen concentrations were averaged and set as representing the trunk neural tube. A two-point calibration was performed using the newly shaken MQ (100% oxygen saturation) and by adding sodium dithionite to non-flowing MQ in the plate after measurements (0% oxygen saturation). Salinity of the tissue was determined using a conductivity meter (WTW 3110) and room temperature noted. The tissue is considered a liquid, where full oxygen saturation at 5 ‰ salinity and 25°C corresponds to 250 μm/l, 160 mmHg or 21% atmospheric O_2_. Data was averaged for each HH stage including one measurement of the previous and subsequent HH stages. Replicates vary from three to ten biologically independent data points. Data is presented as percent of maximum saturation in the solution of the specific temperature and salinity.

### Quantifications

Embryonic development was quantified in two ways; by determining the HH stage of embryos *in ovo* using head and tail morphology or by counting the number of somites of dissected embryos *ex ovo*. The number of embryos (n) for each group is denoted in respective figure legend. The fraction of proliferating EdU+ cells was determined by quantifying the number of GFP+ proliferating cells as well as RFP+ construct targeted cells and divide the number of double positive cells with the number of RFP+ cells. Only neural crest cells were included (distinguished by the dotted line in figures).

### Statistical methods and data sets

One-way ANOVA or two-sided student’s unpaired *t* test was used for statistical analyses. Publicly available dataset Cancer Cell Line Encyclopedia (CCLE) (R2: microarray analysis and visualization platform (http://r2.amc.nl)) was used to analyze gene expression across cell lines from different cancer types. For downstream analysis on the 97 DEGs where the software IPA was used, the statistical tests considered were p-value calculations using right-tailed Fisher Exact Test.

## Acknowledgments

We would like to thank Erica Hutchins, Shashank Gandhi and Siv Beckman for skillful technical assistance and Anni Glud and Ronnie N. Glud for providing microsensor technique and expertise. This work was supported by the Swedish Cancer Society, the Swedish Childhood Cancer Fund, the Craaford Foundation, Jeansson Foundations, Ollie and Elof Ericsson’s foundation, the Mary Bevé Foundation, Magnus Bergvall’s foundation, the Thelma Zoéga foundation for medical research, Hans von Kantzow’s foundation, the Royal Physiographic Society of Lund, the Gyllenstierna Krapperup’s Foundation, and Gunnar Nilssons Cancerstiftelse (to SM), DE027568 and R01HL14058 (to MEB). We thank Center for Translational Genomics, Lund University and Clinical Genomics Lund, SciLifeLab for providing sequencing service. Support by NBIS (National Bioinformatics Infrastructure Sweden) is gratefully acknowledged.

## Author contributions

SM, CUP, EF and EH performed experiments. SM, EH and MEB analyzed data. SM and JML analyzed RNA sequencing data. EM and ZK provided materials. SM and MEB supervised the study. SM wrote the original draft of the manuscript while all authors reviewed and edited the manuscript.

---

**Supplemental Fig. S1.**
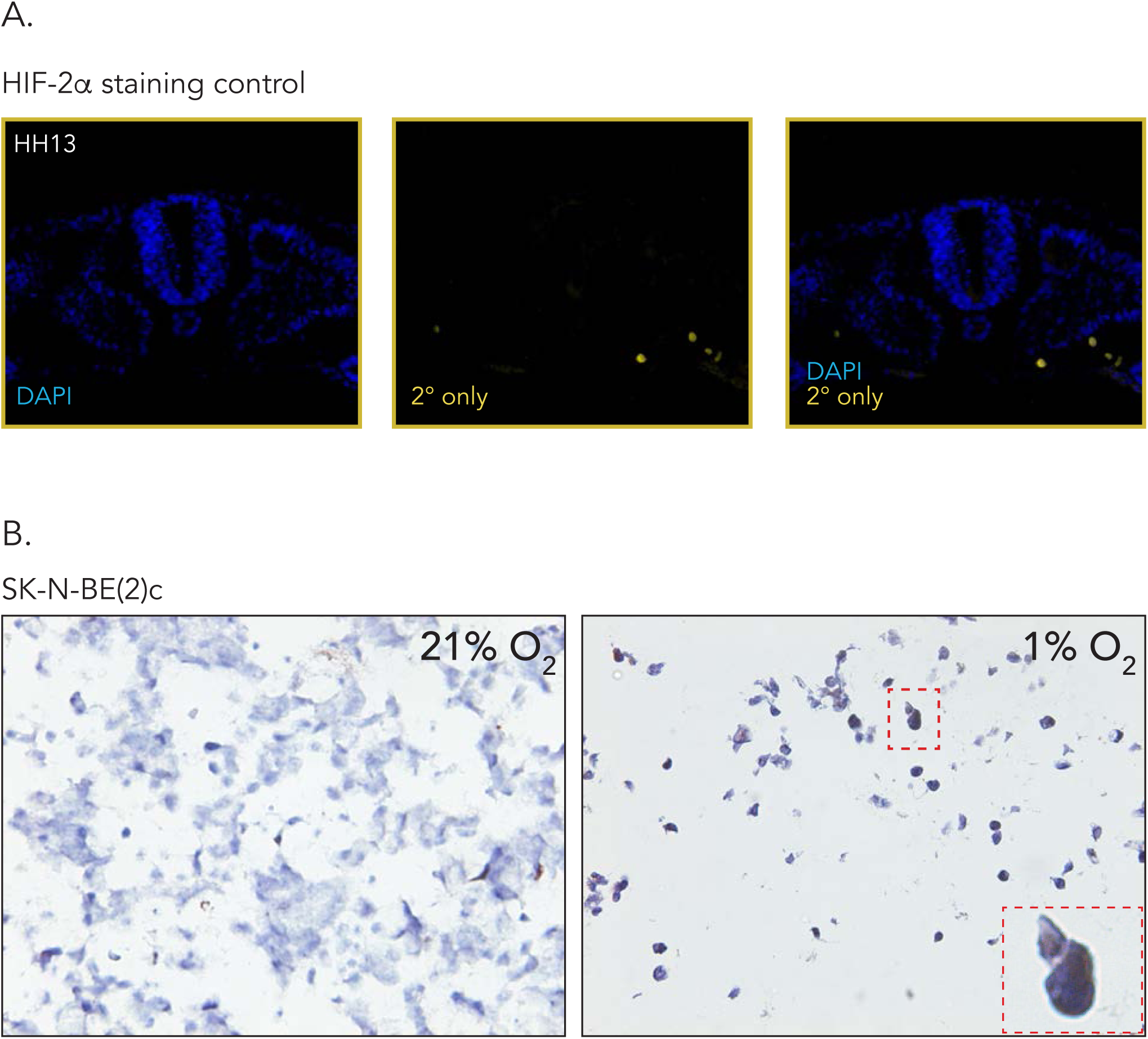
Specificity control of antibodies. **A.** Sections of HH13 wild type embryo immunostained with DAPI for visualization of nuclei and secondary antibody only (donkey anti-rabbit Alexa Fluor-546). **B.** Immunohistochemical staining for HIF-2α in sections of SK-N-BE(2)c neuroblastoma cells cultured at normoxia (21% O_2_) or hypoxia (1% O_2_). HIF-2α positive cells are as expected detected at hypoxia and demonstrate nuclear and cytoplasmic expression.

**Supplemental Fig. S2.**
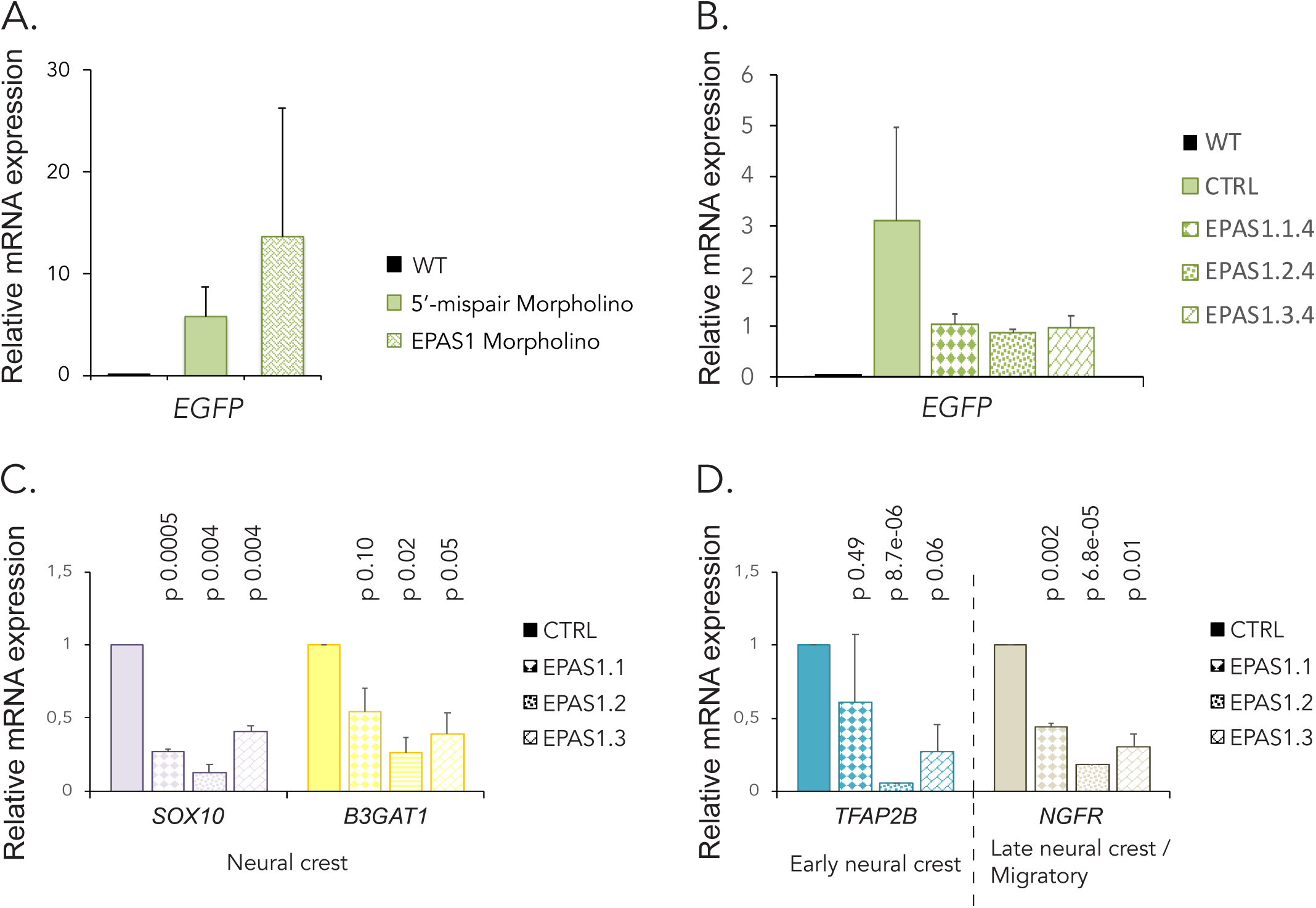
Electroporation of knockdown constructs is efficient. **A-B.** Relative mRNA expression of *EGFP* in embryos electroporated with morpholinos (cf. Figure 3E-F) **(A)** or CRISPR constructs (cf. Figure 3G**-H**) **(B)** measured by qRT-PCR. Expression of *EGFP* in electroporated embryos was compared to expression in wild type HH18 embryos. **C-D.** Neural crest **(C)** and early or late/migratory neural crest **(D)** genes in trunk neural crest cells derived from embryos electroporated with non-targeting (CTRL) or three *EPAS1* gRNAs, measured by qRT-PCR 24 hours post-electroporation. Data are presented as mean of n=2 biologically independent replicates and error bars represent SEM. Statistical significance comparing each individual *EPAS1* targeting gRNA to control (CTRL), respectively, was determined by two-sided student’s t-test.

**Supplemental Fig. S3.**
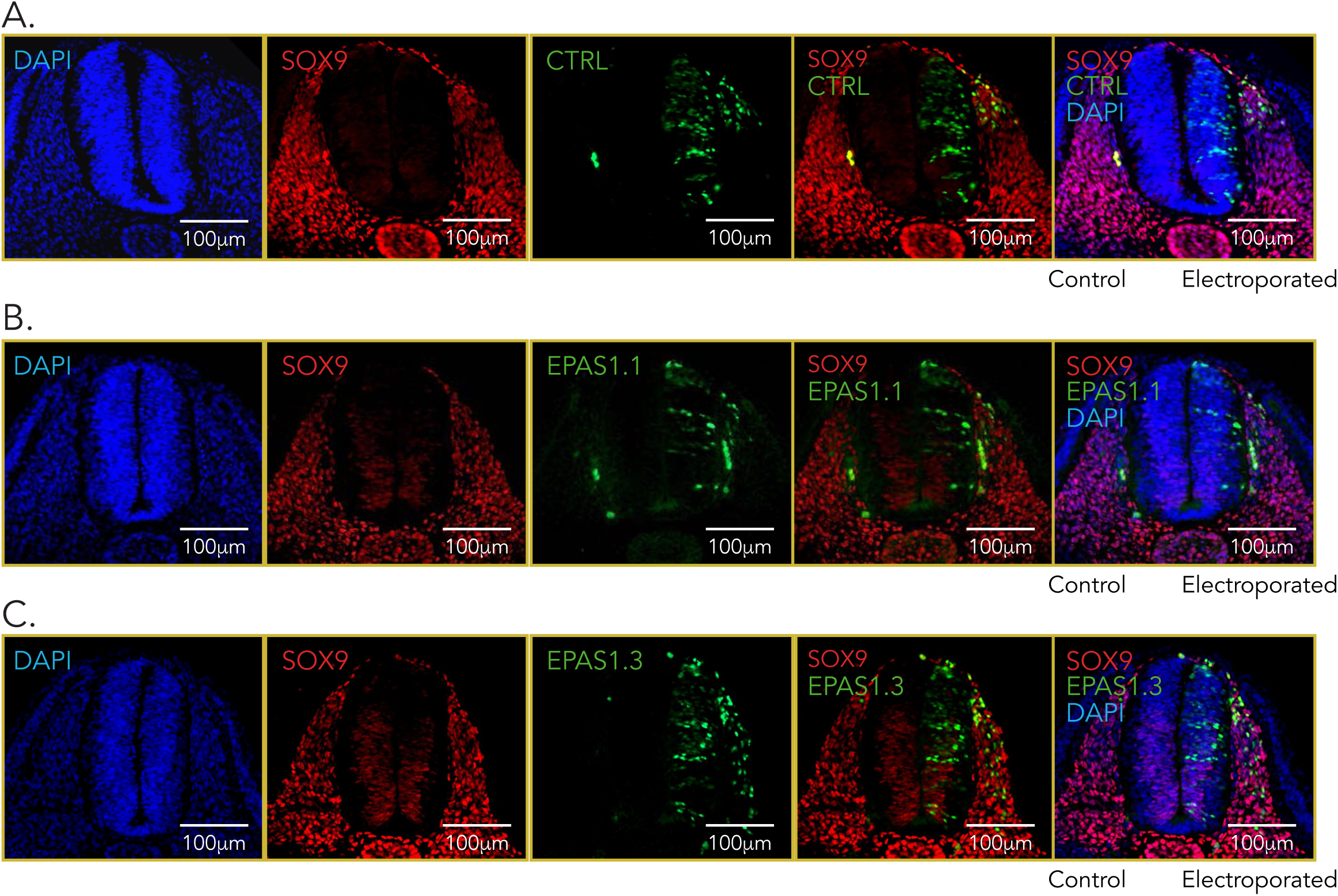
SOX9 expression is not affected by HIF-2α knockdown. **A-C.** Immunostaining of SOX9 (red) in one-sided electroporated embryos (right side). Electroporated cells (non-targeting gRNA (CTRL), **(A)**) or gRNA #1 (EPAS1.1, **(B)**) and #3 (EPAS1.3, **(C)**) targeting EPAS1) are seen in green. DAPI was used to counterstain nuclei. Embryo sections from trunk axial level are from 48 hours post-electroporation.

**Supplemental Fig. S4.**
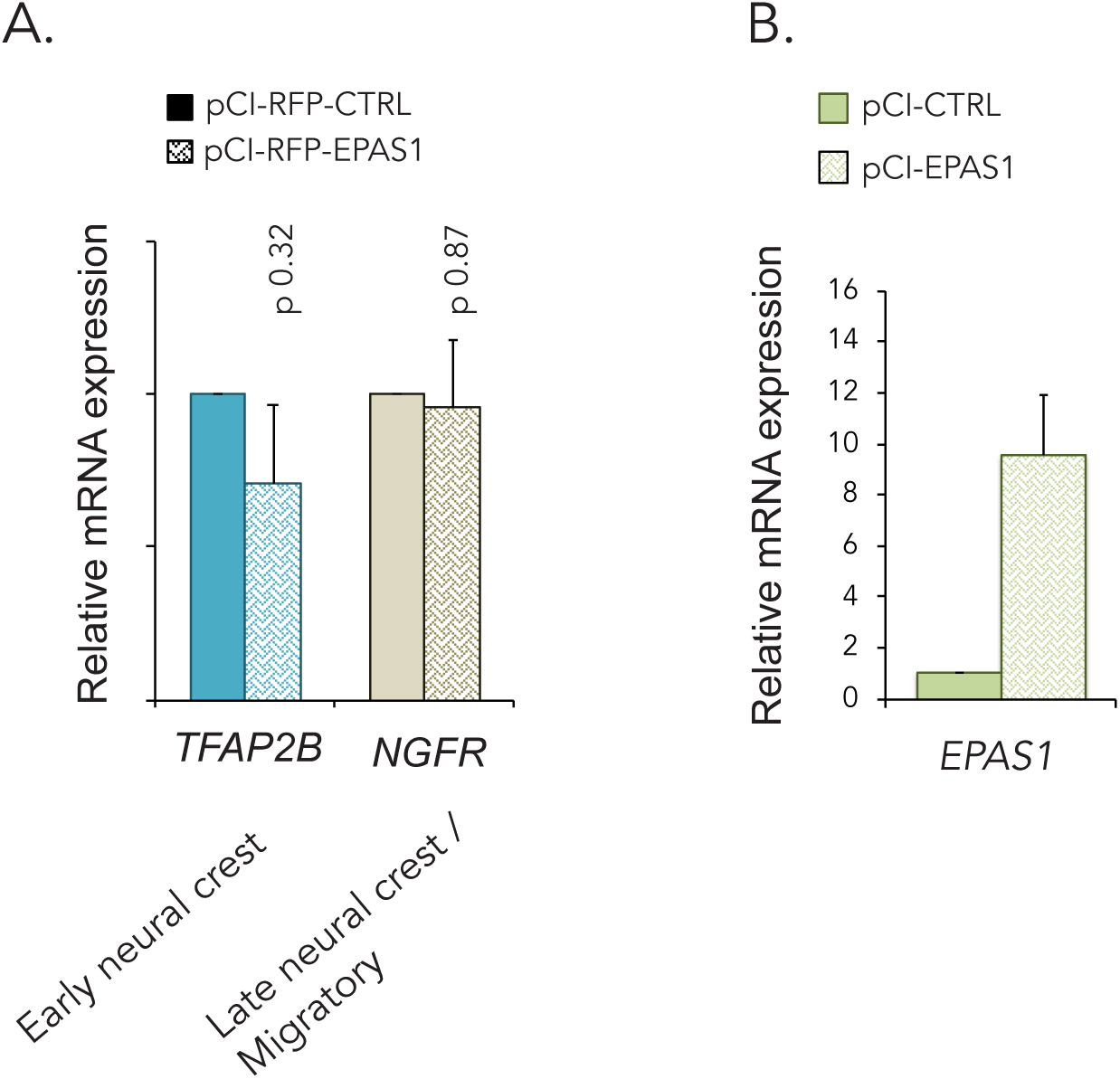
Electroporation of overexpression constructs is efficient. **A.** Relative mRNA expression of neural crest associated genes in trunk neural crest cells derived from embryos electroporated with pCI-CTRL or pCI-EPAS1 vectors, measured by qRT-PCR 24 hours post-electroporation. Data presented as mean of n=2 biologically independent repeats, error bars denote SEM. Statistical significance was determined by two-sided student’s t-test. **B.** Relative mRNA expression of *EPAS1* in embryos electroporated with pCI-CTRL or pCI-EPAS1 for overexpression of HIF-2α (cf. Figure 5). Data are presented as mean of n=2 biologically independent replicates and error bars represent SEM.

**Supplemental Fig. S5.**
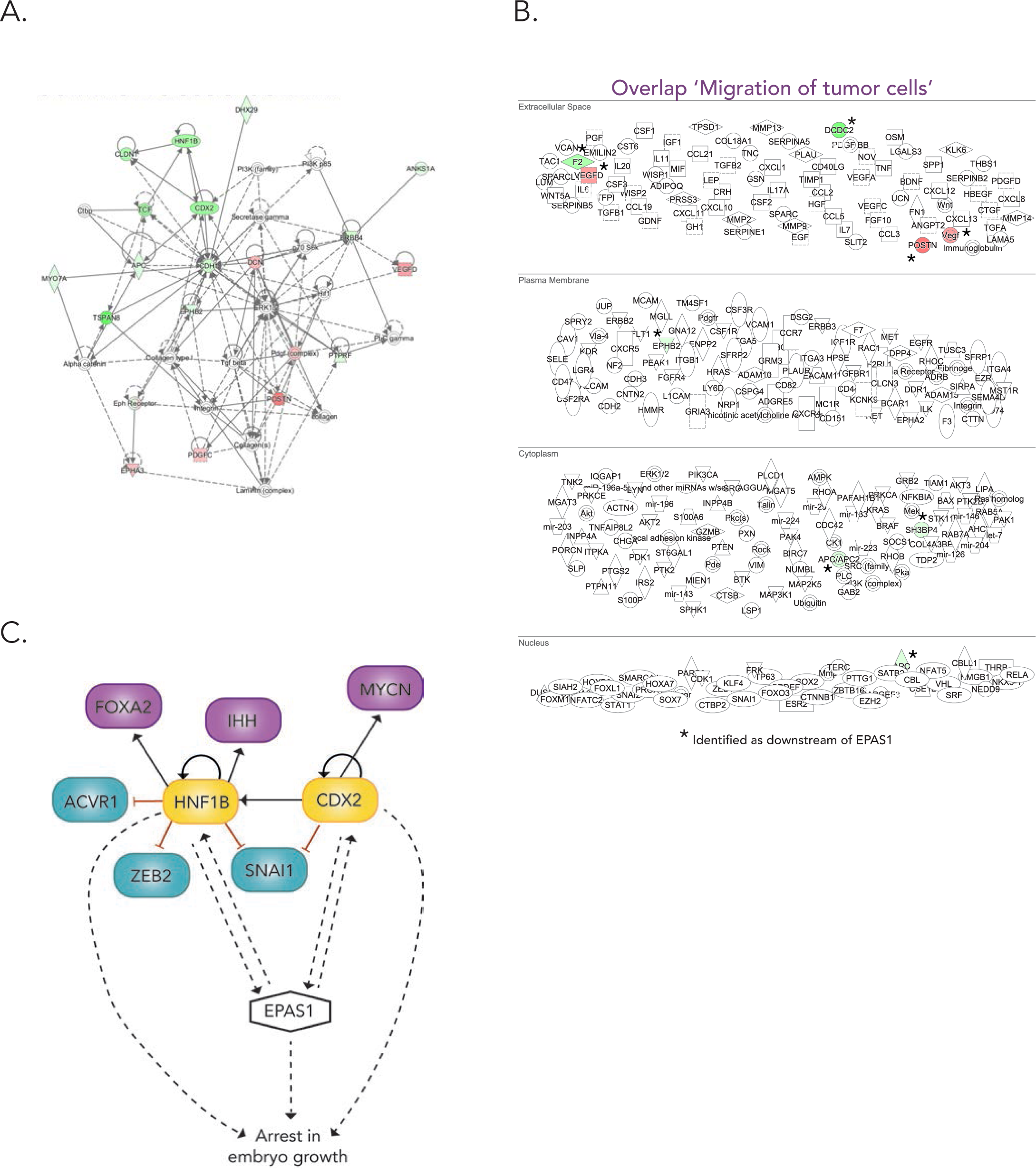
Gene set enrichment analysis identifies key molecules. **A.** Top network composed by analyzing significantly Differentially Expressed Genes from RNA sequencing data. **B.** Deeper analysis of overlap of genes involved in downstream process “migration of tumor cells” and genes from RNA sequencing data. **A-B.** The shape of molecules and their meaning, i.e. correspondence to protein family etc., is found here: http://qiagen.force.com/KnowledgeBase/KnowledgeIPAPage?id=kA41i000000L5rTCAS. As an example, the diamond shaped molecules correspond to enzymes, oval standing shapes should be read as transmembrane receptors and lying oval shapes are transcription regulators. Green nodes indicate down-regulated molecules. The intensity of the color reveals the strength of the expression i.e. the stronger the color the more significant. The dashed lines indicate an indirect interaction between molecules in the network whereas solid lines are direct interactions. The solid arrow explains the direction of the indicated interaction. A line, solid or dashed, without an arrowhead indicate an RNA-RNA interaction. **C.** Schematic of the gene regulatory network including *EPAS1* and downstream *CDX2* and *HNF1B* coupled to arrested embryo growth.

**Supplemental Fig. S6.**
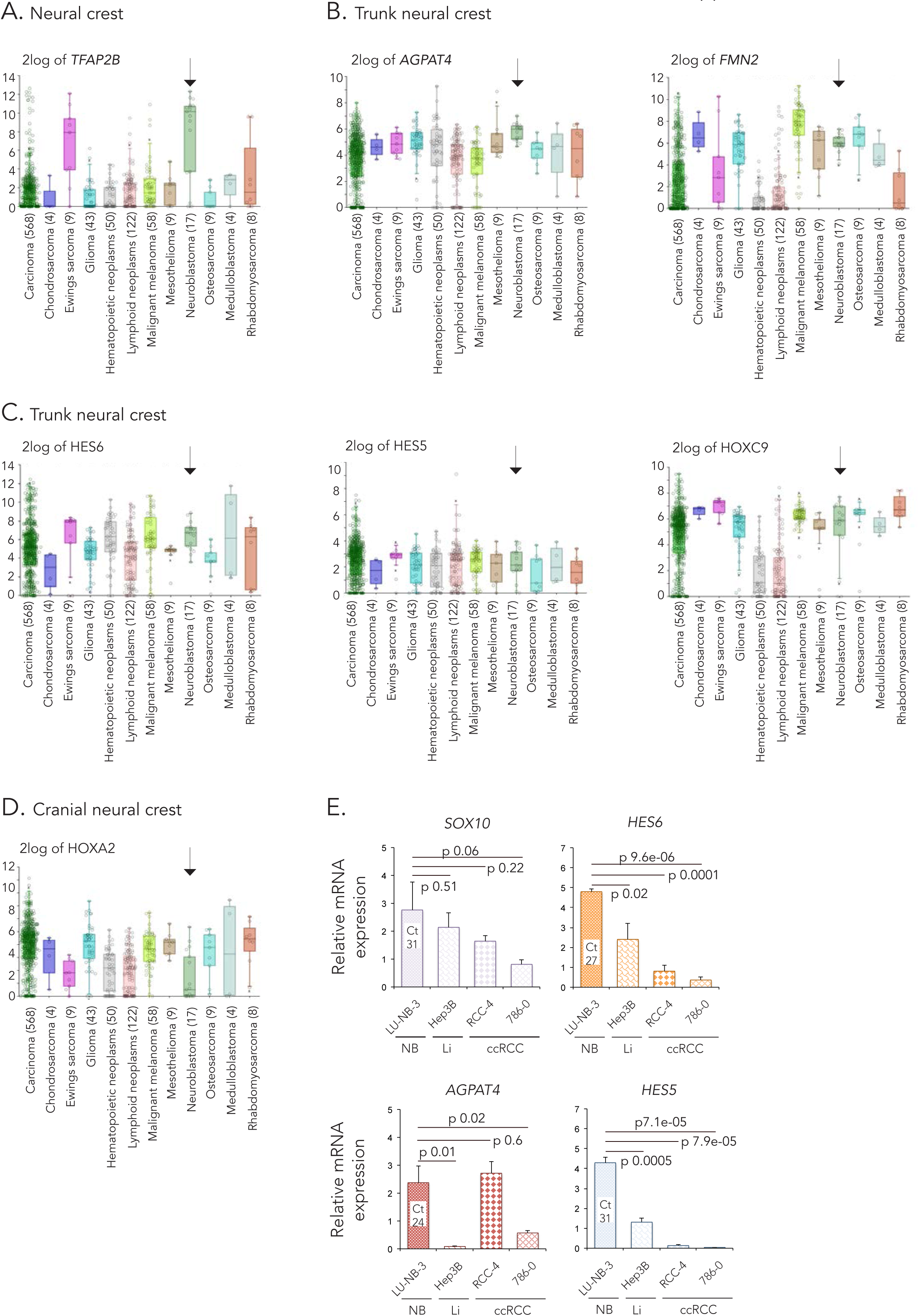
Trunk neural crest associated genes are enriched in neuroblastoma. **A-D.** Neural crest (*TFAP2B* **(A)**), trunk neural crest (*AGPAT4*, *FMN2*, *HES6*, *HES5* and *HOXC9* **(B-C)**) and cranial neural crest (*HOXA2* **(D)**) gene expression in cancer types of different tissue origins. Data from the Cancer Cell Line Encyclopedia (CCLE) dataset, tissue origin with samples n>3 were chosen for further analysis. Arrows highlight neuroblastoma. **E.** Relative mRNA expression of neural crest (*SOX10*) and trunk neural crest (*HES6, AGPAT4, HES5*) genes measured by qRT-PCR. Expression in LU-NB-3 neuroblastoma (NB) patient-derived xenograft cells were compared to liver cancer (Li) Hep3B and clear cell renal cell carcinoma (ccRCC) RCC-4 and 786-0 cell lines. Data are presented as mean of n=3 biologically independent replicates and error bars represent SEM. Statistical significance comparing Hep3B, RCC-4 or 786-0 to LU-NB-3, respectively, was tested using two-sided students *t test*.

**Supplemental Table S1.**
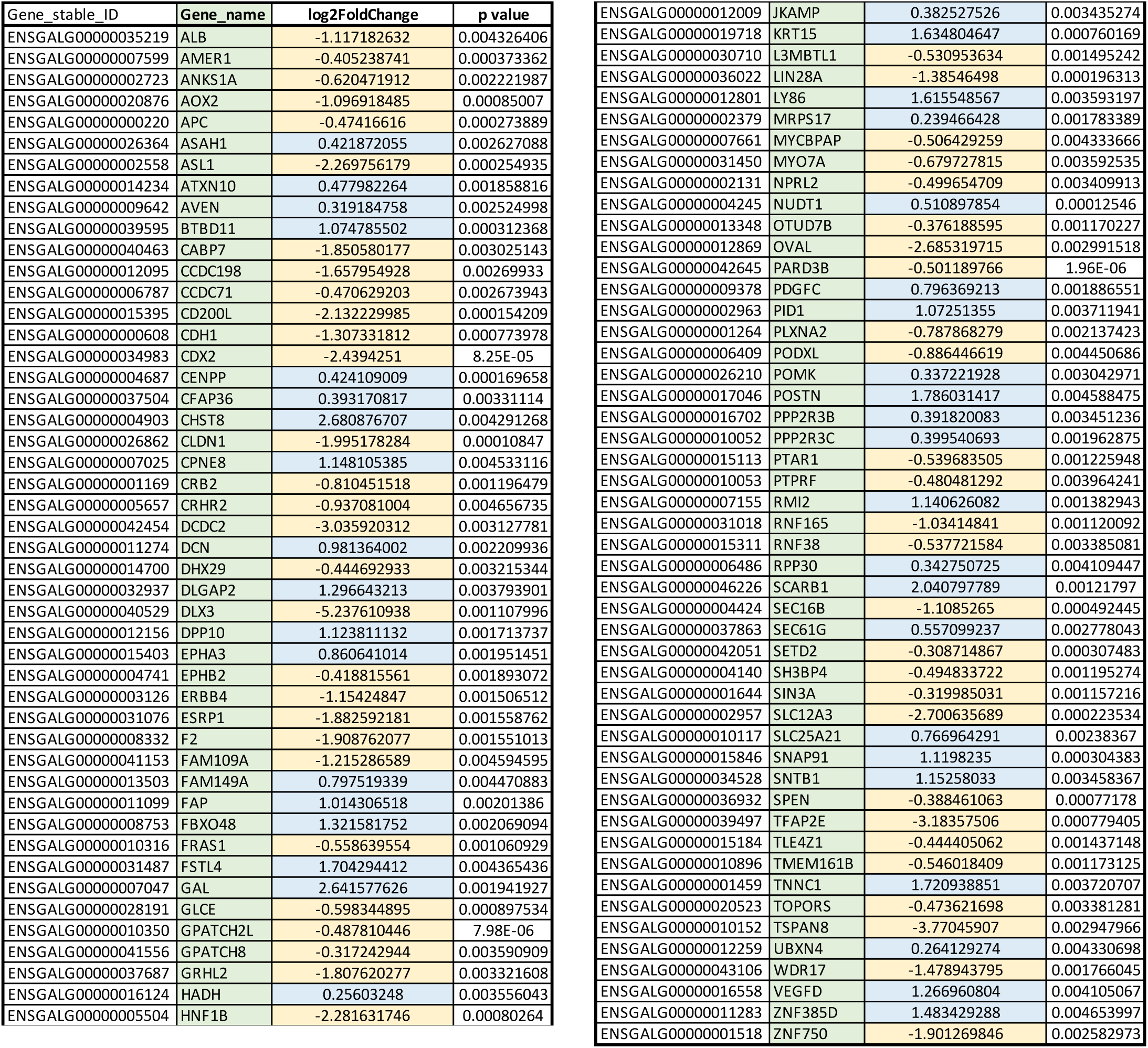
Full list of the 97 significantly (p<0.005) DEGs between 5’-mispair and *EPAS1* morpholino samples identified by RNA sequencing. Relates to Fig. 7A-B.

**Supplemental Table S2.** Full list of processes identified by PANTHER analysis. Relates to Fig. 7F.

|  | Fold Enrichment | p value |
| --- | --- | --- |
| cytolysis by symbiont of host cells (GO:0001897) | > 100 | 1.44E-04 |
| hemolysis in other organism involved in symbiotic interaction (GO:0052331) | > 100 | 1.44E-04 |
| cytolysis in other organism involved in symbiotic interaction (GO:0051801) | > 100 | 2.30E-06 |
| maintenance of mitochondrion location (GO:0051659) | > 100 | 1.44E-04 |
| trans-synaptic signaling by trans-synaptic complex, modulating synaptic transmission (GO:0099557) | > 100 | 1.44E-04 |
| hemolysis in other organism (GO:0044179) | > 100 | 1.44E-04 |
| hemolysis by symbiont of host erythrocytes (GO:0019836) | > 100 | 1.44E-04 |
| killing of cells in other organism involved in symbiotic interaction (GO:0051883) | > 100 | 4.01E-06 |
| disruption of cells of other organism involved in symbiotic interaction (GO:0051818) | > 100 | 4.01E-06 |
| cytolysis in other organism (GO:0051715) | > 100 | 4.01E-06 |
| multi-organism cellular process (GO:0044764) | 60.51 | 3.21E-05 |
| cytolysis (GO:0019835) | 55.01 | 4.07E-05 |
| disruption of cells of other organism (GO:0044364) | 50.43 | 5.06E-05 |
| killing of cells of other organism (GO:0031640) | 50.43 | 5.06E-05 |
| axonal fasciculation (GO:0007413) | 40.34 | 8.99E-05 |
| neuron projection fasciculation (GO:0106030) | 40.34 | 8.99E-05 |
| ephrin receptor signaling pathway (GO:0048013) | 26.03 | 2.58E-05 |
| positive regulation of phosphatidylinositol 3-kinase signaling (GO:0014068) | 17.17 | 1.16E-04 |
| positive regulation of cellular protein localization (GO:1903829) | 7.55 | 4.91E-05 |
| regulation of cellular protein localization (GO:1903827) | 7.09 | 5.64E-07 |
| regulation of cellular localization (GO:0060341) | 4.52 | 1.54E-05 |
| cell migration (GO:0016477) | 4.37 | 2.16E-05 |
| cell motility (GO:0048870) | 3.98 | 1.11E-04 |
| regulation of protein localization (GO:0032880) | 3.94 | 5.84E-05 |
| localization of cell (GO:0051674) | 3.94 | 5.84E-05 |
| locomotion (GO:0040011) | 3.73 | 2.39E-05 |
| regulation of cell differentiation (GO:0045595) | 3.43 | 2.27E-06 |
| regulation of response to stimulus (GO:0048583) | 3.01 | 1.48E-05 |
| regulation of biological process (GO:0050789) | 2.61 | 3.94E-05 |
| regulation of cellular component organization (GO:0051128) | 2.57 | 4.79E-05 |
| regulation of multicellular organismal process (GO:0051239) | 2.55 | 1.41E-05 |
| positive regulation of cellular process (GO:0048522) | 2.33 | 1.37E-05 |
| positive regulation of biological process (GO:0048518) | 2.27 | 1.36E-05 |
| negative regulation of cellular process (GO:0048523) | 2.25 | 7.91E-05 |
| negative regulation of biological process (GO:0048519) | 2.09 | 5.55E-06 |
| cytolysis by symbiont of host cells (GO:0001897) | 2.06 | 2.05E-06 |
| regulation of multicellular organismal development (GO:2000026) | 2.04 | 2.70E-05 |
| regulation of developmental process (GO:0050793) | 2.03 | 8.99E-05 |
| anatomical structure development (GO:0048856) | 2.02 | 6.47E-05 |
| multicellular organism development (GO:0007275) | 1.98 | 3.99E-05 |
| developmental process (GO:0032502) | 1.97 | 6.32E-05 |
| positive regulation of metabolic process (GO:0009893) | 1.85 | 3.35E-05 |
| regulation of metabolic process (GO:0019222) | 1.47 | 1.21E-04 |
| positive regulation of cellular metabolic process (GO:0031325) | > 100 | 1.44E-04 |

**Supplemental Table S3.** Full list of genes identified as potential upstream regulators of “arrest in embryo growth”. Relates to **Table 1**.

| IF antibodies |  |  |  |  |
| --- | --- | --- | --- | --- |
| Primary Antibody | Species | Dilution | Source | Product # |
| HNK1 | Mouse | 1:5 | Hybridoma bank | 3H5 |
| HIF-2α | Rabbit | 1:50 | Abcam | ab199 |
| SOX9 | Rabbit | 1:1000 | Millipore | ab5535 |
| Secondary Antibody | Species | Dilution | Source |  |
| Anti-Mouse Alexa Fluor-594 | Goat | 1:1000 | Invitrogen | A-11032 |
| Anti-Rabbit Alexa Fluor-546 | Donkey | 1:1000 / 1:500 | Invitrogen | A-10040 |
| Anti-Mouse Alexa Fluor-488 | Goat | 1:1000 | Invitrogen | A-11008 |

| IHC antibodies |  |  |  |  |
| --- | --- | --- | --- | --- |
| Primary Antibody | Species | Dilution | Source | Product # |
| HIF-2α | Mouse | 1:1000 | Novus Biologicals | NB100-132 |
| HIF-2α | Rabbit | 1:4000 | Abcam | ab199 |
| TH | Rabbit | 1:1600 | Abcam | ab112 |

| In situ antibodies |  |  |  |  |
| --- | --- | --- | --- | --- |
|  | Species | Dilution | Source | Product # |
| Anti-dig-AP | Mouse | 1:2000 | Roche Diagnostics | 11093274910 |

| Nuclear staining |  |  |  |  |
| --- | --- | --- | --- | --- |
|  | Species | Dilution | Source | Product # |
| DAPI |  | 1:3000 | Dako | D3571 |

| Western blot antibodies |  |  |  |  |
| --- | --- | --- | --- | --- |
| Primary Antibody | Species | Dilution | Source | Product # |
| HIF-2α | Rabbit | 1:200 | Abcam | ab199 |
| SDHA | Mouse | 1:4000 | Abcam | ab14715 |
| Secondary Antibody | Species | Dilution | Source | Product # |
| Anti-Rabbit | Monkey | 1:3000 | Invitrogen | 65-6120 |
| Anti-Mouse | Sheep | 1:5000 | Invitrogen | 62-6520 |

**Supplemental Table S4.**
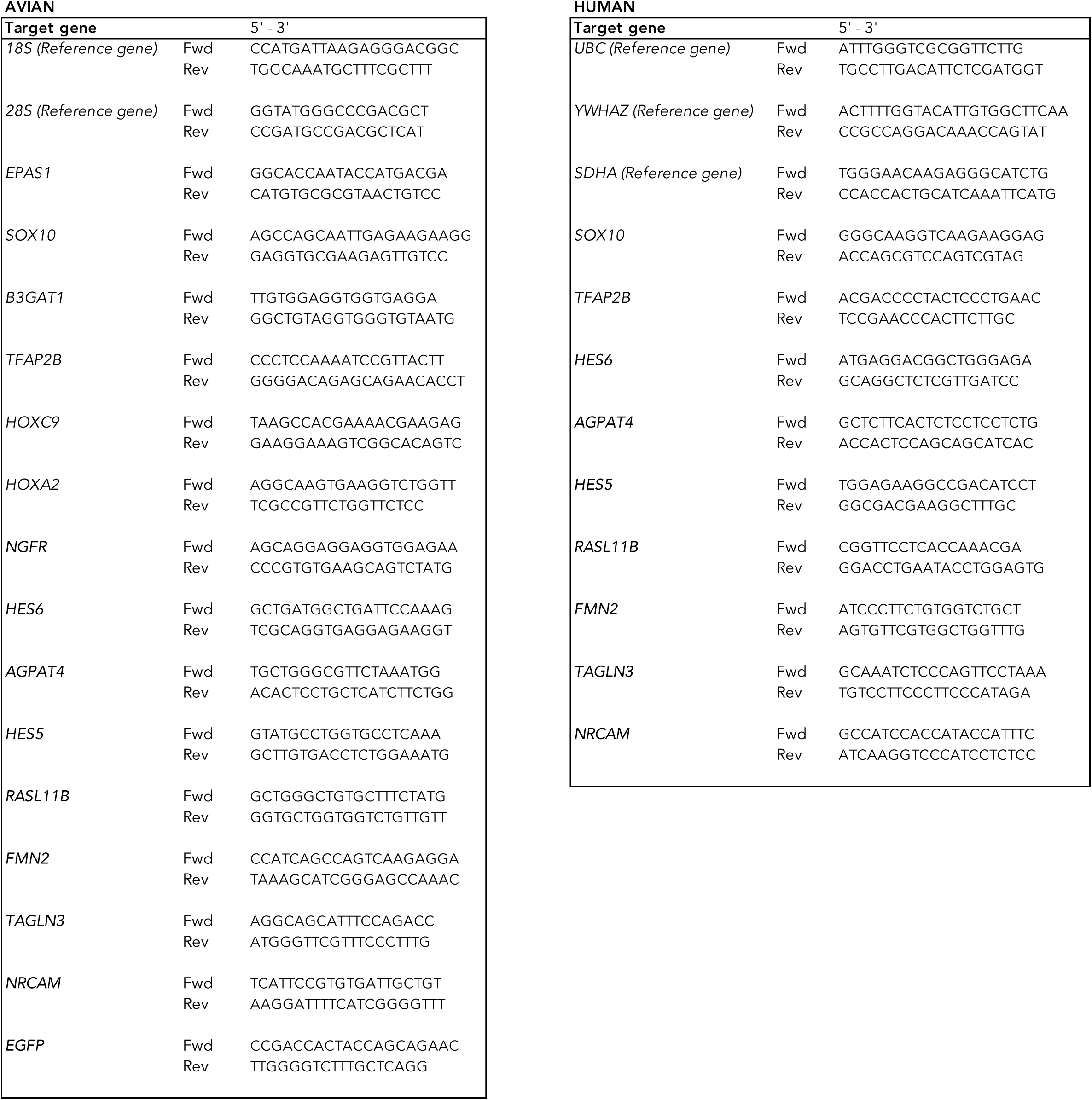
Full list of genes identified as potential upstream regulators of HIF-2α from RNA sequencing data. Target molecules are among the 97 significantly (p<0.005) DEGs between 5’-mispair and *EPAS1* morpholino samples identified by RNA sequencing.

**Supplemental Table S5.** Details of antibodies.

**Supplemental Table S6.** List of primer sequences used for qRT-PCR analyses.

